# Eight new genomes of organohalide-respiring *Dehalococcoides mccartyi* reveal evolutionary trends in reductive dehalogenase enzymes

**DOI:** 10.1101/345173

**Authors:** Olivia Molenda, Shuiquan Tang, Line Lomheim, Elizabeth A. Edwards

## Abstract

**Background:** Bioaugmentation is now a well-established approach for attenuating toxic groundwater and soil contaminants, particularly for chlorinated ethenes and ethanes. The KB-1 and WBC-2 consortia are cultures used for this purpose. These consortia contain organisms belonging to the Dehalococcoidia, including strains of *Dehalococcoides mccartyi* in KB-1 and of both *D. mccartyi* and *Dehalogenimonas* in WBC-2. These tiny anaerobic bacteria couple respiratory reductive dechlorination to growth and harbour multiple reductive dehalogenase genes (*rdhA*) in their genomes, the majority of which have yet to be characterized.

**Results:** Using a combination of Illumina mate-pair and paired-end sequencing we closed the genomes of eight new strains of *Dehalococcoides mccartyi* found in three related KB-1 sub-cultures that were enriched on trichloroethene (TCE), 1,2-dichloroethane (1,2-DCA) and vinyl chloride (VC), bringing the total number of genomes available in NCBI to 24. A pangenome analysis was conducted on 24 *Dehalococcoides* genomes and five *Dehalogenimonas* genomes (2 in draft) currently available in NCBI. This Dehalococcoidia pangenome generated 2875 protein families comprising of 623 core, 2203 accessory, and 49 unique protein families. In *Dehalococcoides mccartyi* the complement of reductive dehalogenase genes varies by strain, but what was most surprising was how the majority of *rdhA* sequences actually exhibit a remarkable degree of synteny across all *D. mccartyi* genomes. Several homologous sequences are also shared with *Dehalogenimonas* genomes. Nucleotide and predicted protein sequences for all reductive dehalogenases were aligned to begin to decode the evolutionary history of reductive dehalogenases in the Dehalococcoidia.

**Conclusions:** The conserved synteny of the *rdhA* genes observed across *Dehalococcoides* genomes indicates that the major differences between strain *rdhA* gene complement has resulted from gene loss rather than recombination. These *rdhA* have a long evolutionary history and trace their origin in the Dehalococcoidia prior to the speciation of *Dehalococcoides* and *Dehalogenimonas.* The only *rdhA* genes suspected to have been acquired by lateral gene transfer are protein-coding *rdhA* that have been identified to catalyze dehalogenation of industrial pollutants. Sequence analysis suggests that evolutionary pressures resulting in new *rdhA* genes involve adaptation of existing dehalogenases to new substrates, mobilization of *rdhA* between genomes or within a genome, and to a lesser degree manipulation of regulatory regions to alter expression.

## 1.0 BACKGROUND

Bioaugmentation using mixed microbial consortia capable of reductive dechlorination is commonly used in attenuating chlorinated aliphatic hydrocarbons (CAHs) in groundwater and soil. Exposure to CAHs is of public concern due to their known toxicity and/or carcinogenicity [1]. The efficacy of *in-situ* bioaugmentation to transform contaminants such as perchloroethene (PCE) and downstream products such as trichloroethene (TCE), *cis-*dichloroethene (cDCE) and vinyl chloride (VC) to ethene is already well established [2, 3]. Reductive dechlorination of these contaminants, especially with two or fewer chlorine constituents, is attributed to the anaerobic bacteria from the class Dehalococcoidia including *Dehalococcoides mccartyi* [4, 5] and *Dehalogenimonas* [6, 7]. The KB-1 mixed microbial consortium is well-suited for this purpose containing multiple strains of *Dehalococcoides mccartyi* capable of complete detoxification of PCE and 1,2-dichloroethane (1,2-DCA) to ethene [8]. WBC-2 contains both *Dehalococcoides* and *Dehalogenimonas* strains that work in concert with other bacteria to dechlorinate a variety of chlorinated ethenes and ethanes. [2, 9].

*Dehalococcoides and Dehalogenimonas* are obligate organohalide respiring bacteria. Reductive dechlorination is carried out by reductive dehalogenase enzymes used to respire CAHs and other halogenated organic compounds and obtain energy for growth. Previous comparisons of *Dehalococcoides* genomes only revealed that they contain a core syntenic region encoding “housekeeping” genes such as biosynthesis of amino acids, cell components, transcription/translation, nutrient transport and energy conservation [10, 11]. The differences between strains in this region are no more than single nucleotide polymorphisms (SNPs). This stable core genome has been described to be interrupted by two variable regions flanking the origin commonly referred to as High Plasticity Regions (HPR1 and HPR2) [10, 11]. HPRs show signs of recombination including repeats, duplication events, insertion sequences, genomic islands, phage related sequences and hold the majority of reductive dehalogenases which are used for organohalide respiration. The primary type of recombination event observable in the HPRs is site-specific recombination - involving the reciprocal exchange of DNA between defined DNA sites [12]. *D. mccartyi* genomes contain as many as ten recombinases per genome in contrast to *Escherichia coli* K-12 which has six, despite a genome that is ~3.5 times larger. Prokaryotes also use recombination to change gene expression by manipulating regulatory sequences relative to coding sequences [13, 14], or while accepting DNA from conjugation [15]. Several characterized dehalogenases lie on genomic islands with evidence of site-specific integration, including the VC reductase genes *vcrA* and *bvcA* [16] and the TCE reductase gene *tceA* [17]. It is thought that recombination events have allowed *D. mccartyi* to adapt to naturally-occurring and anthropogenic halogenated compounds [11, 16].

*D. mccartyi* and *Dehalogenimonas* diverged from their most recent Dehalococcoidia ancestor anywhere from 40,000 to 400,000 years ago [16, 18]. Dehalococcoidia are present in both contaminated and uncontaminated environments playing a role in the global halogen cycle unrelated to releases of organohalides from anthropogenic activities [19]. These bacteria have small genomes (~1.4 Mbp for *D. mccartyi*, ~1.7 Mbp for *Dehalogenimonas*), all *Dehalococcoides* sequenced to date are from the same species (*mccartyi*), while sequenced *Dehalogenimonas* span multiple species. Circumstances which favour a small genome can be explained by the genome streamlining hypothesis [20-22]. Small genome sizes could also be caused as a by-product of niche specialization due to increased rates of mutation [23] or as a result of gene loss [24]. The evolution of genomes in general is thought to be dominated by long term reduction and simplification, with brief episodes of complexification in response to environmental conditions [25]. Evidently, *D. mccartyi* and *Dehalogenimonas* must have experienced long term reduction and simplification to achieve their current state. Some reductive dehalogenase genes have very few mutations at the nucleotide or amino acid level suggesting that they have not been a part of *D. mccartyi* genomes for very long. Over time, a gene under strong selective pressure will acquire more synonymous mutations than non-synonymous as a result of purifying selection. The *vcrA* gene encoding for the vinyl chloride reductase has very few mutations, being 98% conserved across eight different strains of *D. mccartyi*, suggesting that it was recently horizontally distributed across populations possibly in response to industrial activities and the release of chlorinated ethenes into the environment [16].

Reductive dehalogenase genes typically occur in an operon containing the gene for the reductive dehalogenase catalytic A subunit (*rdhA*) and membrane anchor (*rdhB*) and sometimes other genes. On an amino acid basis, the overall similarity between dehalogenases can be very low [26] and classification is usually based on the presence of three motifs: a TAT export sequence, two FeS clusters, and a corrinoid-binding domain. While many dehalogenase sequences can be identified in any particular *D. mccartyi* genome, only a select few have been found expressed in response to different CAHs [27, 28]. As a result, only a few dehalogenases have been characterized out of the over five hundred Dehalococcoidia sequences that are found in ref_seq in NCBI. These include VcrA (primarily VC, but also TCE, DCE and 1,2-DCA to ethene [29]), TceA (primarily TCE to cDCE [30]), PceA (PCE to TCE and 2,3-dichlorophenol (2,3-DCP) to monochlorophenol [28]), BvcA (cDCE, VC and 1,2-DCA to ethene [31]), MbrA (PCE to tDCE [32]), CbrA (1,2,3,4-tetrachlorobenzene to 1,2,4-trichlorobenzene, also 1,2,3- trichlorobenzene to 1,3-dichlorobenezene [33]), PteA (TCE to ethene [34]),TdrA (tDCE to VC [35]), DcpA (1,2-DCA to ethene [36]) and CerA (VC to ethene [6]). Organohalide respiring bacteria are widely distributed among different phylogenies [37], and several additional dehalogenases have been characterized from outside the Dehalococcoidia. The only structures available are from a respiratory reductive dehalogenase, PceA, from *Sulfurosprillum multivorans* [38] and a catabolic (intracellular) reductive dehalogenase RdhANP from *Nitratireducter pacificus* [39]. Hug *et al.* [18] proposed a naming system for grouping dehalogenase sequences for ease of comparison, despite most not having ascribed substrate. A relatively arbitrary cut off of 90% amino acid similarity was proposed to define sets of orthologous RdhA sequences (i.e. Ortholog Groups, or OG) which likely share the same or similar substrates [18, 35].

Several strains of *D. mccartyi*, including the type strain 195 [40], GT [41] and BAV1 [42] have been obtained in pure culture. No isolates from the KB-1 culture have been obtained despite multiple attempts. Nevertheless, Pérez-de-Mora *et al.* [43] deduced from quantitative PCR analyses that multiple distinct strains of *D. mccartyi* were present in KB-1 sub-cultures even after years of enrichment on a single chlorinated substrate. The objectives of this study were to identify and characterize the different strains of *D. mccartyi* found in KB-1 subcultures amended with different chlorinated electron acceptors. From the metagenomes of these cultures, 8 new *D. mccartyi* genomes were closed, increasing the total number of publicly available genomes to 24. These genomes were compared within the *Dehalococcoides* and with other available *Dehalogenimonas* sequences to build a pangenome and to begin to identify factors influencing genome evolution, DNA recombination and specifically in the evolution of dehalogenase genes. DNA and amino acid sequence information was used to particularly investigate reductive dehalogenases that have not yet been biochemically characterized.

## 2.0 METHODS

### 2.1 Enrichment cultures analysed

The KB-1 cultures were originally enriched from aquifer materials at a TCE contaminated site in southern Ontario as previous described [2, 3, 42]. Four different enrichment cultures were created. The parent culture was first enriched in 1996 with TCE and methanol as electron donor (KB-1/TCE-MeOH). A subculture was created in 2001 that was amended with VC and H_2_ as electron donor (KB-1/VC-H_2_ culture [42]). In 2003, two additional subcultures of the parent culture were established, one with cDCE and methanol (KB-1/cDCE-MeOH), and one with 1,2-DCA and methanol (KB-1/1,2-DCA-MeOH). These cultures have been maintained on these substrates ever since; details of culture maintenance are provided in supplemental information.

### 2.2 Metagenomic sequencing and genome assembly

DNA for metagenome sequencing was extracted from larger samples (40-615 mL) taken from the three stable enrichment cultures described above: KB-1/VC-H_2_ (40 mL culture sample), KB-1/TCE-MeOH (500 mL sample), KB-1/cDCE-MeOH (300 mL culture) and KB-1/1, 2-DCA-MeOH (615 mL sample). Extractions were conducted between February and May, 2013. Cultures were filtered using Sterivex™ filters (Millipore 0.2 µm) and the DNA was extracted using the CTAB method (JGI bacterial genomic DNA isolation using CTAB protocol v.3). DNA was sequenced at the Genome Quebec Innovation Sequencing Centre using Illumina HiSeq 2500 technology. Paired-end sequencing with an insert size of ~400 bp and read length of ~150 bp provided roughly 50 million reads per culture. Additional mate-pair sequencing with insert size of ~8000 bp and read length of ~100 bp was conducted for the KB-1/TCE-MeOH and KB-1/1, 2- DCA-MeOH cultures where we had more DNA. In the case of metagenomic sequencing using short-read Next Generation Sequencing (NGS), we have demonstrated the utility of long-insert paired-end data in resolving challenges in metagenomic assembly, especially those related to repeat elements and strain variation [44]. In this study, we applied Illumina mate-pair sequencing to incorporate such data. Although other long-read sequencing technologies (e.g. PacBio SMRT sequencing, Nanopore sequencing, Illumina Synthetic Long-Read Sequencing Technique and 10x Genomics) are also used, Illumina mate-pair sequencing is a cost-effective choice for the goal of obtaining both high sequencing depth and accuracy and long-distance mate pair links. Raw sequences were trimmed with Trimmomatic [45] to remove bases of low quality and to remove adapters.

The *D. mccartyi* genomes were assembled in six steps as described below. In <u>Step 1</u> we generated ABySS unitigs with Illumina Paired-end data using ABySS assembler [46]. These unitigs were the main building blocks in the assembly of the complete genomes. ABySS assemblies generate both unitigs and contigs. Unlike contigs, unitigs are generated solely by overlapping k-mers and their assembly does not utilize the paired-end constraints. As a result, the maximum overlapping length between unitigs is the length of k-mer size minus one. When using the ABySS to assemble metagenomic data, we used the maximum k-mer size allowed, 96 bp, since the raw read length was 150bp, much longer than 96 bp. When configuring ABySS runs, it was critical to utilize the -c parameter, which specifies a cut-off, the minimum k-mer depth/coverage used in the assembly. Sequences/unitigs with k-mer coverage lower than this cutoff will be ignored in the assembly, which allows users to have good assemblies of high abundance organisms as the interferences caused by low abundance organisms (especially those of close relatives) and by sequencing errors are removed. It is important to make sure that the k-mer depth of the sequences of the target genomes is higher than this threshold so that you have all sequences/unitigs you need to close the target genomes. For example, if the average k-mer depth of the target genome is 100, try 20 for the -c cut off. We used a combination of 16S rRNA amplicon sequencing and qPCR to get an idea of the relative abundances of our target organisms in our metagenome prior to attempting different ABySS assemblies.

In <u>Step 2</u> we generated a genome-wide reference sequence for the target genome, which will be subsequently be used to guide the scaffolding of unitigs. This reference sequence can be obtained in different ways. If there are long-distance mate-paired data as what we had for KB-1/T3MP1-MeOH and KB-1/1, 2-DCA-MeOH cultures, this reference sequence can be built de novo. We used two ways to build it: (1) using a standalone scaffolding program, SSPACE v. 2.0 [47], to generates scaffolds with ABySS contigs/unitigs utilizing the mate-paired constraints, (2) using ALLPATHS-LG [48] to generate the assembly with both paired-end and mate-pair data as inputs. ALLPATHS-LG turned out to be the most effective way in most cases. A publicly available closely related closed genome might also be able to serve as a reference genome to guide scaffolding of unitigs in the next step.

One major challenge in metagenomic assembly is cross-interference between closely related genomes, such as strains of the same species. The sequence similarity/dissimilarity between these closely related genomes tend to break the assembly. If a genome had a closely related genome interfering in its assembly, we attempted to assemble the genome with ALLPATHS-LG using both short-insert paired-end data and long-insert mate-pair data. For genomes that have no closely related genomes, a surprisingly effective way to assemble is to combine Digital Normalization [49] with ALLPATHS-LG. This approach reduces the data redundancy of raw sequences with Digital Normalization by k-mer and then one can assemble the resulting data with ALLPATHS-LG. In our case, we had multiple *D. mccartyi* strains in each metagenome and could not use Digital Normalization. ALLPATHS-LG was able to differentiate our similar strains because their abundances were distinct.

In <u>Step 3,</u> we used the best assembly generated from Step 2 to guide the scaffolding of unitigs generated from Step One. The scaffolding process is based on sequence comparison between the unitigs and the reference assembly by BLAST. After that, unitigs that have a k-mer depth significantly lower or higher than the average k-mer depth of the genome are removed. The basic assumption here is that unitigs with k-mer depth higher than average likely belong to repetitive sequences (such as rRNA gene operons and transposons) and unitigs with a k-mer depth lower than average are more likely to be strain specific. In other words, only unitigs with k-mer depth around average (we used 90%-110% of average) are kept; these unitigs are likely shared by closely related genomes. After that, the gap distance between the neighbouring unitigs is estimated based on the reference assembly. In brief, this process generated a scaffold consisting of unitigs shared by all closely related strains; this will service as a backbone for subsequent gap resolution.

In <u>Step 4,</u> we identify all potential solutions for all gaps between unitigs in the scaffold. This step is performed by filling the gaps with the remaining unitigs mostly based on sequence overlap between unitigs; we published a similar process previously [44]. In the updated script (available in supplemental information), we have improved the process by incorporating paired-end and mate-pair link information between unitigs to help guide the searching process. The paired-end and mate-pair links were obtained by mapping raw reads against unitigs. Solutions identified this way fulfill the constraints of sequence overlap, paired-end links and mate-pair links. If there are multiple solutions to a gap and they have k-mer depth lower than the average, this suggests the presence of strain variation. In the end, this step generates a closed assembly, having some gaps with multiple solutions in cases of strain variation.

In <u>Step 5</u> we bin these multiple solutions caused by strain variation to different genomes based on sequencing depth or k-mer depth. For example, if there are always two solutions, one of k-mer depth of 60 and the other one of 40, we will assign all solutions with higher depth to one strain and the rest to the other strain. This approach is unfeasible if the two strains happen to have similar abundance and thus similar sequencing depth. Things become more complicated when there are more than two strains; in such case, we only try to resolve the genome of the highest abundance strains by gathering solutions of highest k-mer depth. The editing of the genome sequences is facilitated by the use of Geneious v. 6.1 [50]. Finally, in <u>Step 6</u> we polish the assembled genome by mapping raw reads back to the final assembly. SNPs caused by strain variation are identified. If possible, they are resolved based on abundance in the same principle as using k-mer depth to assign alternative solutions in Step 5.

In all cases multiple genomes could be closed from a single enrichment culture because the different populations of *D. mccartyi* were at different abundances (as inferred from read depth) at the time of sampling. Two complete genomes each containing a vinyl chloride reductase gene (*vcrA*) were closed from the KB-1/VC-H_2_. The naming convention used here distinguishes KB-1 lineage (KB-1) electron acceptor (in this case vinyl chloride or VC) and relative abundance (number 1 for highest abundance and so on) naming the strains from KB-1/VC-H_2_ culture *D. mccartyi* strains KBVC1 and KBVC2. Three genomes each containing *bvcA* were closed from the KB-1/1, 2-DCA enrichment culture further referred to as strain KBDCA1, KBDCA2 and KBDCA3. Two genomes each containing *tceA* from KB-1/TCE-MeOH culture, strains KBTCE1 and KBTCE2. A *D. mccartyi* complete genome containing *vcrA* gene was also assembled from KB-1/TCE-MeOH culture, strain KBTCE1. In all cases low abundance strains of *D. mccartyi* could not be assembled implying that although eight genomes were closed, the total number of KB-1 *D. mccartyi* strains is at least eleven. Each genome was annotated first using RAST [51], with hypothetical annotations resubmitted to BASys [52] some of which were assigned an annotation. Microbial Genomes Check from NCBI assisted in finding errors produced from automatic annotation. Results were manually inspected and corrected where required using Geneious ORF finder. NCBI automatically annotates reference sequence version using PGAP [53]. Additional searches for conserved domains were conducted using NCBI conserved domain search (E-value threshold of 0.01). The origin of replication was identified using Oriloc in R [54].

### 2.3 Amplicon Sequencing and Analysis

For microbial community analysis, amplicon sequencing was performed on extracted DNA, which was amplified by PCR using general primers for the 16S rRNA gene. The universal primer set, 926f (5’-AAACTYAAAKGAATTGACGG-3’) and 1392r (5’- ACGGGCGGTGTGTRC-3’), targeting the V6-V8 variable region of the 16S rRNA gene from bacteria and Archaea, as well as the 18S rRNA gene from Eukaryota, were used [55]. The purified PCR products were sent to the McGill University and Genome Quebec Innovation Centre, where they were checked for quality again, pooled and subject to unidirectional sequencing (*i.e.*, Lib-l chemistry) of the 16S gene libraries, using the Roche GS FLX Titanium technology (Roche Diagnostics Corporation, Indianapolis, IN). One to three independent 100 µL PCR amplification reactions were preformed per sample. Each PCR reaction was set up in sterile Ultra-Pure H_2_O containing 50uL of PCR mix (Thermo Fisher Scientific, Waltham, MA), 2 µL of each primer (forward and reverse, each from 10 μM stock solutions), and 4 μL of DNA extract. PCR reactions were run on a MJ Research PTC-200 Peltier Thermal Cycler (Bio-Rad Laboratories, Hercules, CA) with the following thermocycling program; 95 °C, 3 min; 25 cycles of 95 °C 30 s, 54 °C 45 s, 72 °C 90 s; 72 °C 10 min; final hold at 4 °C (modified from [56]). The forward and reverse primers included adaptors (926f: CCATCTCATCCCTGCGTGTCTCCGACTCAG and, 1392r: CCTATCCCCTGTGTGCCTTGGCAGTCTCAG), and the reverse primer also included 10bp multiplex identifiers (MID) for distinguishing multiple samples pooled within one sequencing region. The PCR products were verified on a 2% agarose gel and replicates were combined and purified using GeneJET™ PCR Purification Kit (Fermentas, Burlington, ON), according to the manufacturer’s instructions. The concentrations of PCR products were determined using a NanoDrop ND-1000 Spectrophotometer at a wavelength of 260 nm (NanoDrop Technologies, Wilmington, DE). The concentrations and qualities of the final PCR products were also evaluated by running them on 2% agarose gels, and comparing band intensities to those from a serial dilution of ladders with known DNA concentrations.

### 2.4 Taxonomic assignments of 16S rRNA reads

The raw DNA sequences obtained from the sequencing center were processed using the Quantitative Insights Into Microbial Ecology (QIIME v1.5.0) pipeline [57] with default settings, unless stated otherwise. Only sequences of length between 300 and 500 bp, and with homopolymers shorter than 8 bases were processed for downstream analysis. After filtering, sequences were de-multiplexed into respective samples based on their individual MID. Sequences were further clustered into distinct 16S rRNA gene-based Operational Taxonomic Units (OTUs) using the UCLUST algorithm [58], similarity threshold of 0.97 and the Green Genes database (version 13.5) [59]. Taxonomy was assigned to each OTU by the Ribosomal Database Project (RDP) classifier [60].

### 2.5 Quantitative PCR (qPCR) Analysis

qPCR was used to estimate the abundance of specific *rdhA* and *D. mccartyi* sequences in each of the sequenced KB-1 cultures. DNA samples were diluted 10, 50 or 100 times with sterile UV treated distilled water (UltraPure), and all subsequent sample manipulations were conducted in a PCR cabinet (ESCO Technologies, Gatboro, PA). Each qPCR reaction was run in duplicate. Four *Dehalococcoides* genes were targeted by qPCR: 1) the phylogenetic 16S rRNA gene for *Dehalococcoides* Dhc1f (5’-GATGAACGCTAGCGGCG-3’) and Dhc264r (5’-CCTCTCAGACCAGCTACCGATCGAA-3’) [61]; 2) the vinyl chloride reductase gene, *vcrA*, vcrA642f (5’-GAAAGCTCAGCCGATGACTC-3’) and vcrA846r (5’-TGGTTGAGGTAGGGTGAAGG-3’) [62]; 3) *bvcA* dehalogenase, bvcA318f (5’-ATTTAGCGTGGGCAAAACAG-3’) and bvcA555r (5’-CCTTCCCACCTTGGGTATTT-3’) [62]; and 4) *tceA* dehalogenase: tceA500f (5’TAATATATGCCGCCACGAATGG-3’) and tceA795r(5’-AATCGTATACCAAGGCCCGAGG-3’) [28]. Samples were also analysed using general bacteria 16S rRNA primers GenBac1055f (5’-ATGGCTGTCGTCAGCT-3’) and GenBac1392r (5’-ACGGGCGGTGTGTAC-3’) [63]. DNA samples were diluted 10, 50 or 100 times with sterile UltraPure distilled water, and all subsequent sample manipulations were conducted in a PCR cabinet (ESCO Technologies, Gatboro, PA). Each qPCR reaction was run in duplicate. Each qPCR run was calibrated by constructing a standard curve using known concentrations of plasmid DNA containing the gene insert of interest. The standard curve was run with 8 concentrations, ranging from 10 to 10^8^ gene copies/µL. All qPCR analyses were conducted using a CFX96 real-time PCR detection system, with a C1000 thermo cycler (Bio-Rad Laboratories, Hercules, CA). Each 20 µL qPCR reaction was prepared in sterile UltraPure distilled water containing 10 µL of EvaGreen^®^ Supermix (Bio-Rad Laboratories, Hercules, CA), 0.5 µL of each primer (forward and reverse, each from 10 μM stock solutions), and 2 μL of diluted template (DNA extract or standard plasmids). The thermocycling program was as follows: initial denaturation at 95°C for 2 min, followed by 40 cycles of denaturation at 98°C for 5s, annealing at 60°C (for 16S rRNA and *vcrA*, *bvcA* genes, respectively) or 58 °C for *tceA* or 55 oC for General Bacteria followed by extension for 10s at 72 °C. A final melting curve analysis was conducted at the end of the program. R^2^ values were 0.99 or greater and efficiency values 80-110%.

### 2.6 Alignments and phylogenetic trees

A core gene alignment was created by aligning a set of 109 core genes found in *Dehalococcoides mccartyi* (22 strains), *Dehalogenimonas lykanthroporepellens*, *D. alkenigignens, Dehalogenimonas* sp. WBC-2 and a Chloroflexi out-group *Sphaerobacter thermophilus.* Core genes are defined as orthologous genes which are present in all genomes analyzed. Core genes were identified using reciprocal BLASTp followed by manual inspection. Each gene was aligned using muscle v. 3.8.3.1 [64] with default settings. The alignments were concatenated to create one long alignment (138,334 bp long, 26 sequences, 83% pairwise identity). A maximum likelihood (ML) tree was built using RAxML [65] plugin in Geneious 8.1.8 [50] with GTR gamma nucleotide substitution model and 100 bootstrap replicates. The best scoring ML tree was chosen as the final tree.

Five-hundred and fifty one *rdhA* sequences were selected to create a nucleotide phylogenetic tree using Geneious 8.1.8. These included all *rdhA* which have been assigned an ortholog group (OG) number from the RDase database, all *rdhA* from three *Dehalogenimonas (lykanthroporepellens, alkenigignens* and sp. WBC-2) and a reductive dehalogenase from *Desulfoluna spongiiphila* as the out-group. *D. spongiiphila* is an anaerobic, sulfate-reducing bacterium isolated from a marine sponge [66]. The alignment and tree building was conducted using muscle and RAxML as described above. FigTree 1.4.2 was used to visualize and further edit the tree to generate figures in this paper (http://tree.bio.ed.ac.uk/software/figtree/).

The ratio of synonymous substitutions (Ks) to non-synonymous (Ka) was compared. The Ka/Ks ratio can be used to identify positive selection. If all non-synonymous mutations are either neutral or deleterious, then Ka/Ks < 1, while if Ka/Ks > 1, then positive selection occurred [67]. Ka/Ks ratios were calculated using http://services.cbu.uib.no/tools/kaks.

### 2.7 Homologous gene clustering and pangenome analysis

Twenty-four *D. mccartyi* genomes and five *Dehalogenimonas* genomes (2 draft) (KB-1 and from NCBI) were analyzed using the GET_HOMOLOGUES [68] open sourced software package designed for pangenome analysis. The prokaryotic genome pipeline was used to cluster homologous protein families using first BLASTp (min coverage 75 and E-value 1e^-5^) to calculate bidirectional best-hit (BDBH) followed by Markov clustering referred to as OrthoMCL method. Minimum cluster size was one protein sequence. A pangenome matrix was created summarizing which Dehalococcoidia genome had which protein cluster present in its genome, and how many representatives from that cluster.

In order to investigate synteny a series of whole genome alignments was produced using Mauve [69] plugin in Geneious 8.1.8 [50]. Subsequently MCScanX [70] package was chosen to calculate collinear blocks of coding sequences and create figures. All KB-1 *Dehalococcoides mccartyi* genome coding sequences as well as *Dehalogenimonas* WBC-2 (*Dehalococcoidia*) and *Sphaerobacter thermophilus* (a Chloroflexi) coding sequences were compared using BLASTp. One best alignment was chosen for each coding sequence from each of the genomes with an E-value of at least 1e^-2^. MCScanX used BLASTp input to calculate collinear blocks and progressively align multiple collinear blocks between genomes on default settings. MCScanX’s circle plotter program was used to generate figures.

### 2.8 Statistical analysis of pangenome homologous protein clusters

Pearson’s chi-squared test was used to determine whether the proteins found in each homologous cluster differed significantly between *Dehalococcoides mccartyi* and *Dehalogenimonas.* A correspondence analysis (CA) was used to compare the contents of the protein clusters with the genomes they were generated from. In this analysis a total of 2875 homologous protein clusters were generated. A scree plot was used to compare the percentage contributions of each dimension to the expected value (3.6%, if all dimensions contributed equally) in order to only consider significant dimensions. As a result, we reduced the number of dimensions from 28 to 9. The contributions of individual genomes and protein clusters were identified by creating a bar plot of top contributors and comparing to a reference line corresponding to the expected value if the contributions were uniform. Any row/column with a contribution above the reference line was considered important to the final ordination. All analyses were conducted using R. 3.4.0. using FactoMineR, factoextra and vegan packages.

## 3.0 Nucleotide sequence accession numbers

KB-1 *Dehalococcoides mccartyi* closed genome nucleotide accession numbers in the National Center for Biotechnology Information (NCBI): strain KBDCA1 CP019867, strain KBDCA2 CP019868, strain KBDCA3 CP019946, strain KBVC1 CP019968, strain KBVC2 CP19969, strain KBTCE1 CP01999, strain KBTCE2 CP019865, and strain KBCTCE3 CP019866.

KB-1 16S rRNA amplicon sequences have been deposited in NCBI in the short-read archive (SRA) accession no. SRP144609 as part of bioproject no. PRJNA376155.

WBC-2 16S rRNA amplicon are also in NCBI (SRA) accession no. SRP051778 as part of bioproject no. PRJNA269960.

RdhA google database and orthologous groups (OGs) can be retrieved at: https://drive.google.com/drive/folders/0BwCzK8wzlz8ON1o2Z3FTbHFPYXc

## 4.0 RESULTS and DISCUSSION

### 3.1 Microbial diversity in KB-1 and WBC-2 enrichment cultures identified from 16S rRNA amplicon sequencing

The microbial diversity found in the KB-1 consortium and derived sub-cultures has been studied since its first enrichment from TCE-contaminated soils in 1996 [8, 71]. In this study we used 16S rRNA amplicon sequencing to confirm community composition of four different cultures each amended with the same amount of electron donor (5x methanol or 5x hydrogen gas) and different chlorinated substrates including: TCE, cDCE, VC and 1,2-DCA. The main roles of both dechlorinating and non-dechlorinating organisms have been well established in the KB-1/TCE-MeOH enrichment culture with *D. mccartyi* being responsible for dechlorination of TCE and all daughter products to ethene, and *Geobacter* sp. capable of stepwise dechlorination of PCE to cDCE [27, 71]. Non-dechlorinating organisms such as acetogens/fermenters (*Acetobacterium*, Spirochaetaceae, Synergistales), methanogens (*Methanoregula and Methanomethylovorans*) and Firmicutes (*Sporomusa*) degrade methanol to hydrogen or methane and some provide key nutrients such as corrinoid cofactors [71] (Figure 1). While individual organisms carrying out a particular function have been known to vary in relative abundance [72], community level functioning remains consistent. Different techniques have been previously used to track microbial diversity within KB-1 cultures (qPCR, metagenome sequencing [71], and shotgun metagenomic microarray [73]) revealing stability of the microbial community over many decades, also reflected in results presented here (Figure 1, Table S1). The WBC-2 trans-DCE (tDCE) enrichment culture grown with lactate and EtOH as electron donors similarly has fermenting organisms (primarily Veillonellaceae, Spirochaetaceae, Bacteriodales, Figure 1), methanogens (*Methanosphaerula*, Figure 1), *Dehalogenimonas* sp. WBC-2 degrading tDCE to VC [7] and *Dehalococcoides mccartyi* WBC-2 degrading primarily VC to ethene [7, 74]. Veillonellaceae and *Dehalococcoides* were identified in WBC-2 clone libraries as early as 2006 [75], with Bacteriodales in 2007 [76]. The WBC-2 community has also remained stable over seven years of laboratory cultivation.

**Figure 1.**
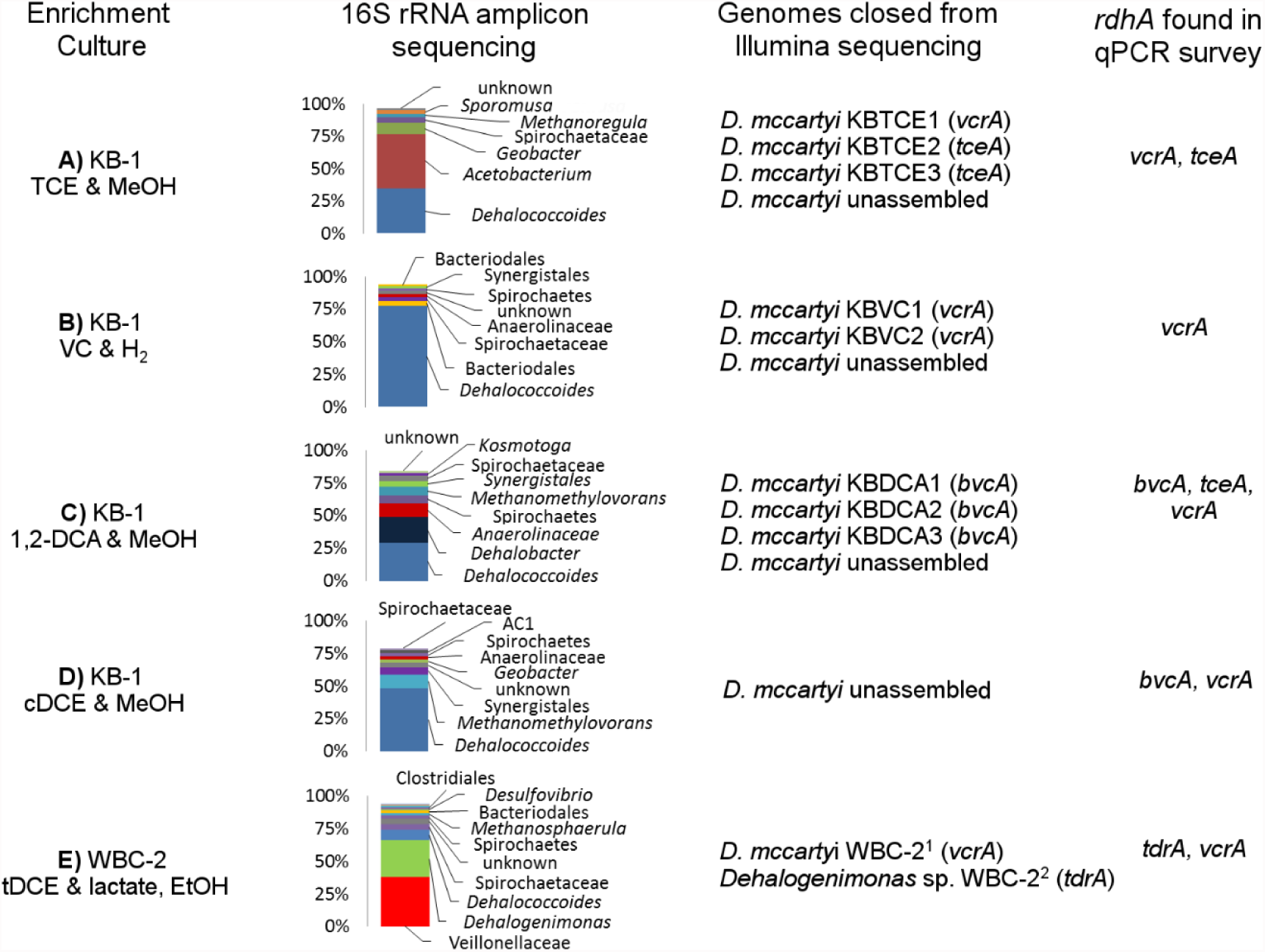
Culture composition and Dehalococcoides mccartyi (Dhc) genomes closed from 16S rRNA amplicon sequencing, Illumina sequencing and qPCR of rdhA genes. DNA was extracted from KB-1 & WBC-2 enrichment cultures. The same DNA sample was split and analysed using 16S rRNA amplicon sequencing (bar charts), Illumina paired-end (all) and mate-pair (all except cDCE enrichment) assembly and genomes closed (genomes listed by rank abundance and strain name if closed) and surveyed for presence of *rdhA* genes using qPCR (final column). Genes found above the detection limit are listed. See supplemental information excel file Table S1 for tabular qPCR results. The eight closed KB-1 *D. mccartyi* genomes shown are from this study. The WBC-2 genomes were previously published in ^1^Genome Announc. 2016 4(6):e01375-16 and ^2^Appl Environ Microbiol. 2015 82(1):40-50.

The main purpose of this round of 16S rRNA amplicon sequencing was to assist with the assembly of mate-pair and paired-end sequencing conducted in order to close Dehalococcoidia genomes. Better assemblies produced longer contigs which aided in genome closing. Figure 1 shows the combined data from 16S rRNA amplicons sequencing, metagenomic Illumina sequencing and qPCR used in concert to close *D. mccartyi* genomes.

### 3.2 General features of new *D. mccartyi* KB-1 genomes

Eight complete genomes of *D. mccartyi* strains were assembled (Figure 2) and annotated from three different enrichment cultures (Table 1) using Illumina mate-pair and paired-end metagenomic sequencing in combination with 16SrRNA amplicon sequencing and qPCR of function *rdhA* to guide assembly (Figure 1). The strains have been named based on contaminant/electron acceptor amended and a number to indicate rank abundance with respect to other strains found in that same enrichment culture (Figure 3). *D. mccartyi* in general have 98% sequence similarity across all 16S rRNA genes [77] (Figure S1) and fall into three clades known as the Pinellas, Victoria and Cornell [78]. Six of the KB-1 strains (KBVC1, KBVC2, KBDCA1, KBDCA2, KBDCA3 and KBTCE1) fall into the Pinellas clade, along with strains CBDB1 [10], BTF08 [79], DCMB5 [79], 11a5 [80], WBC-2 [35], GT [41], IBARAKI [81] and BAV1. Two of the KB-1 strains (KBTCE2 & KBTCE3) fall into the Cornell clade containing strain 195, MB [82] and CG4 [83]) (Figure S1). All new genomes have a clear GC skew with one origin of replication, similar to previously described *Dehalococcoides* genomes. The majority of reductive dehalogenase genes continue to be found in HPRs flanking the origin of replication, primarily coded on opposite leading strands.

**Table 1.** General features of *Dehalococcoides mccartyi* genomes closed from KB-1 trichloroethene (TCE), 1,2-dichloroethane (1,2-DCA) and vinyl chloride (VC) enrichment cultures compared to type strain 195.

|  | KBVC1 | KBVC2 | KBTCE1 | KBTCE2 | KBTCE3 | KBDCA1 | KBDCA2 | KBDCA3 | 195<br>(Reference) |
| --- | --- | --- | --- | --- | --- | --- | --- | --- | --- |
| Genome size (Mbp) | 1.39 | 1.35 | 1.39 | 1.33 | 1.27 | 1.43 | 1.39 | 1.34 | 1.47 |
| G+C content (%) | 47.3 | 47.2 | 47.3 | 49.1 | 49.3 | 47.4 | 47.5 | 47.6 | 48.9 |
| Protein coding genes | 1468 | 1432 | 1451 | 1381 | 1319 | 1496 | 1462 | 1404 | 1582 |
| Hypothetical genes (%) | 31.1 | 30.2 | 30.3 | 29.1 | 26.8 | 32.8 | 31.9 | 29.1 | 34.4 |
| tRNA | 47 | 48 | 47 | 45 | 45 | 47 | 46 | 46 | 46 |
| CRISPR-Cas genes | 7 | 0 | 0 | 0 | 0 | 0 | 0 | 6 | 0 |
| Nitrogen fixation genes | 0 | 0 | 0 | 9 | 9 | 0 | 0 | 0 | 10 |
| Serine recombinases | 2 | 0 | 0 | 5 | 2 | 5 | 4 | 2 | 2 |
| Tyrosine recombinases | 2 | 4 | 4 | 5 | 5 | 5 | 5 | 2 | 2 |
| Sub-group/Clade | Pinellas | Pinellas | Pinellas | Cornell | Cornell | Pinellas | Pinellas | Pinellas | Cornell |
| Electron Acceptor<br>provided to Culture | VC | VC | TCE | TCE | TCE | 1,2-DCA | 1,2-DCA | 1,2-DCA | PCE |
| *rdhA* genes | 22 | 16 | 16 | 5 | 5 | 7 | 7 | 9 | 18 |
| Identifiable *rdhA* * | VcrA,<br>PceA | VcrA,<br>PceA | VcrA,<br>PceA | TceA | TceA | BvcA | BvcA | BvcA | TceA,<br>PceA |
| NCBI accession<br>number | CP019968 | CP019969 | CP019999 | CP019865 | CP019866 | CP019867 | CP019868 | CP019946 | NC002936 |

\* Identifiable *rdhA* indicates presence of *rdhA* gene whose protein product was characterized in a different study. It is not known whether these are currently expressed and by these strains.

**Figure 2.**
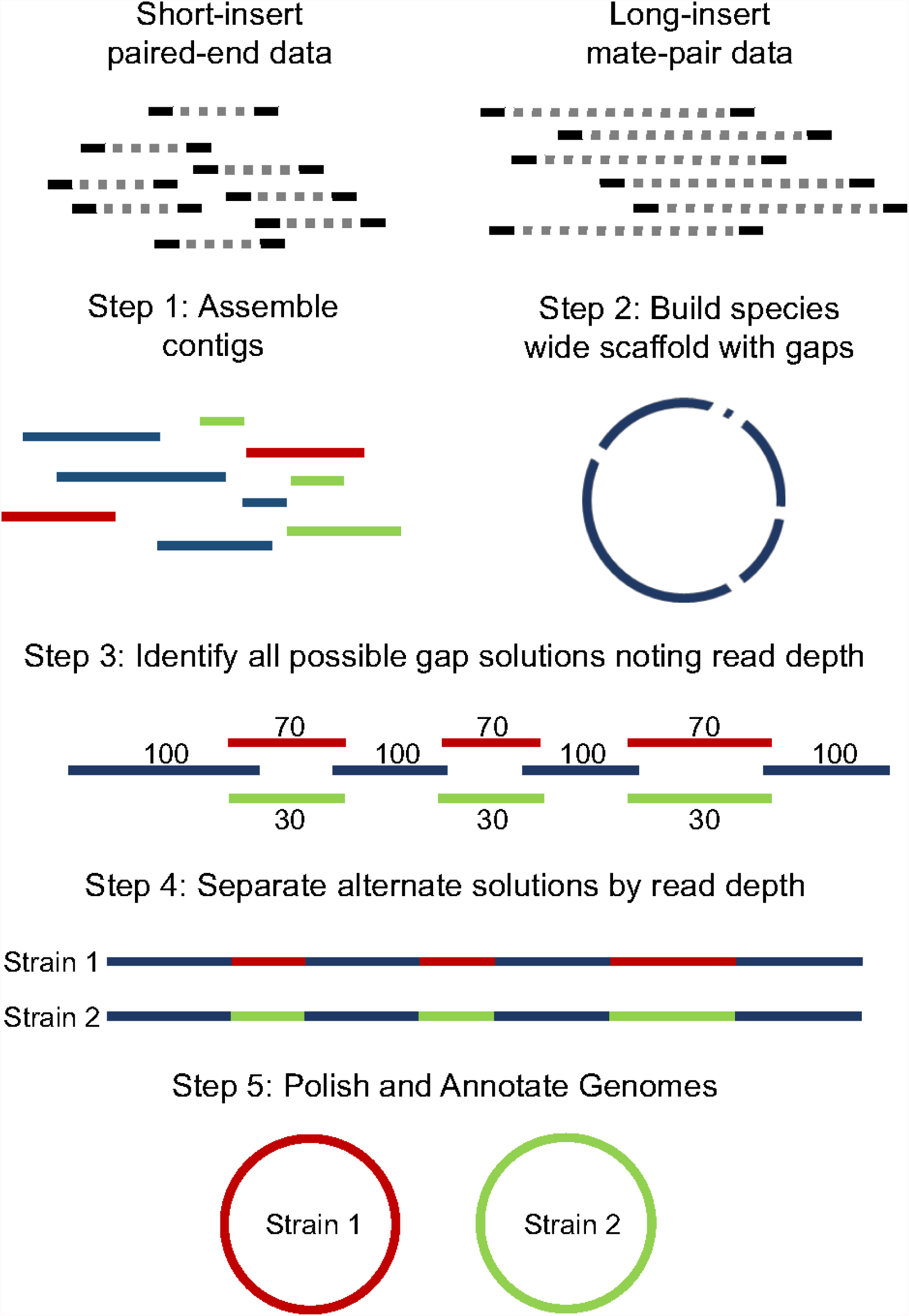
Schematic flow chart of workflow used to assemble *Dehalococcoides mccartyi* Genomes from KB-1 metagenomes.

**Figure 3.**
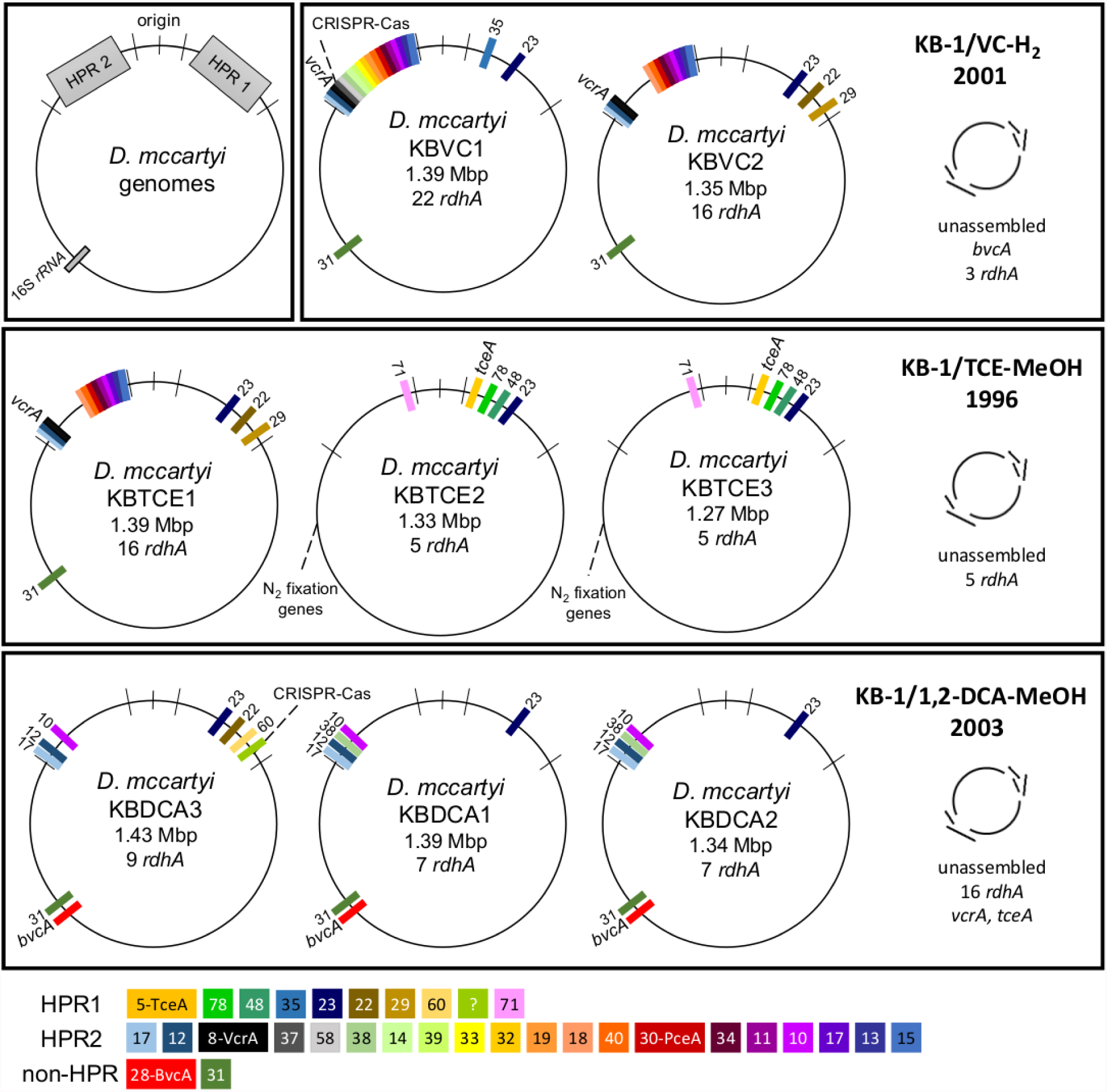
Overview of *D. mccartyi* genomes closed from three different KB-1 enrichment cultures. Each culture is labeled by electron acceptor and electron donor and the date the culture was first created. Trichloroethene (TCE), vinyl chloride (VC), and 1,2-dichloroethane (1,2-DCA) were the electron acceptors for each enrichment culture. Genomes which could be closed are identified by a name indicating electron acceptor and rank abundance as determined from read depth. Reductive dehalogenase homologous genes (*rdhA*) are marked on each genome, coloured by orthologous (OG) group. HPR- High plasticity regions HPR1 and HPR2.

Strains KBVC1 and KBDCA3 each contain a complete CRISPR-Cas system [84] as do published strains 11a [85], CBDB1 [10], DCBM5 [79] and GT [41]. Strains KBTCE2 and KBTCE3 contain nitrogen fixation genes as does strain 195 [86]. The number of putative reductive dehalogenase genes (*rdhA*) varies from five to twenty-two per KB-1 *Dehalococcoides* genome. In general, these new genomes have some of the features which have already been seen in other *D. mccartyi* strains. However, we did find the smallest *D. mccartyi* genome closed to date (strain KBTCE3) at 1.27 Mbp with highest GC content (49.3%) containing the fewest *rdhA* sequences (only 5) of any *D. mccartyi* genome to date (Table 1).

### 3.3 Reductive dehalogenase genes in *Dehalococcoides mccartyi* and conserved synteny

Hug *et al.* (2013) [18] developed a classification system for reductive dehalogenases where sequences were assigned to orthologous groups (OGs) based on ≥ 90% amino-acid similarity to attempt to cluster *rdhA* sequences into groups with activity on the same specific halogenated electron acceptors [35] (Figure 4). The database is available on Google drive (https://drive.google.com/drive/folders/0BwCzK8wzlz8ON1o2Z3FTbHFPYXc) and is user-updated to include new strains. The database was designed to encompass all respiratory reductive dehalogenase homologs, in this paper we focus on Dehalococcoidia RdhA specifically. Previous studies identified 31 distinct *rdhA* genes in the KB-1/TCE-MeOH culture [43, 71]. Although hundreds of putative RdhA sequences were found in KB-1 metagenomes, only two new OG could be described. The remaining *rdhA* fell into already identified groupings (Figure 4). In general, as more *D. mccartyi* genomes are closed, fewer dehalogenases are found that don’t already belong to an OG. (Figure S2) At this time there are 84 distinct OGs and 31 RdhA sequences which cannot be grouped on the basis of >90% amino acid similarity, and remain as singletons.

**Figure 4.**
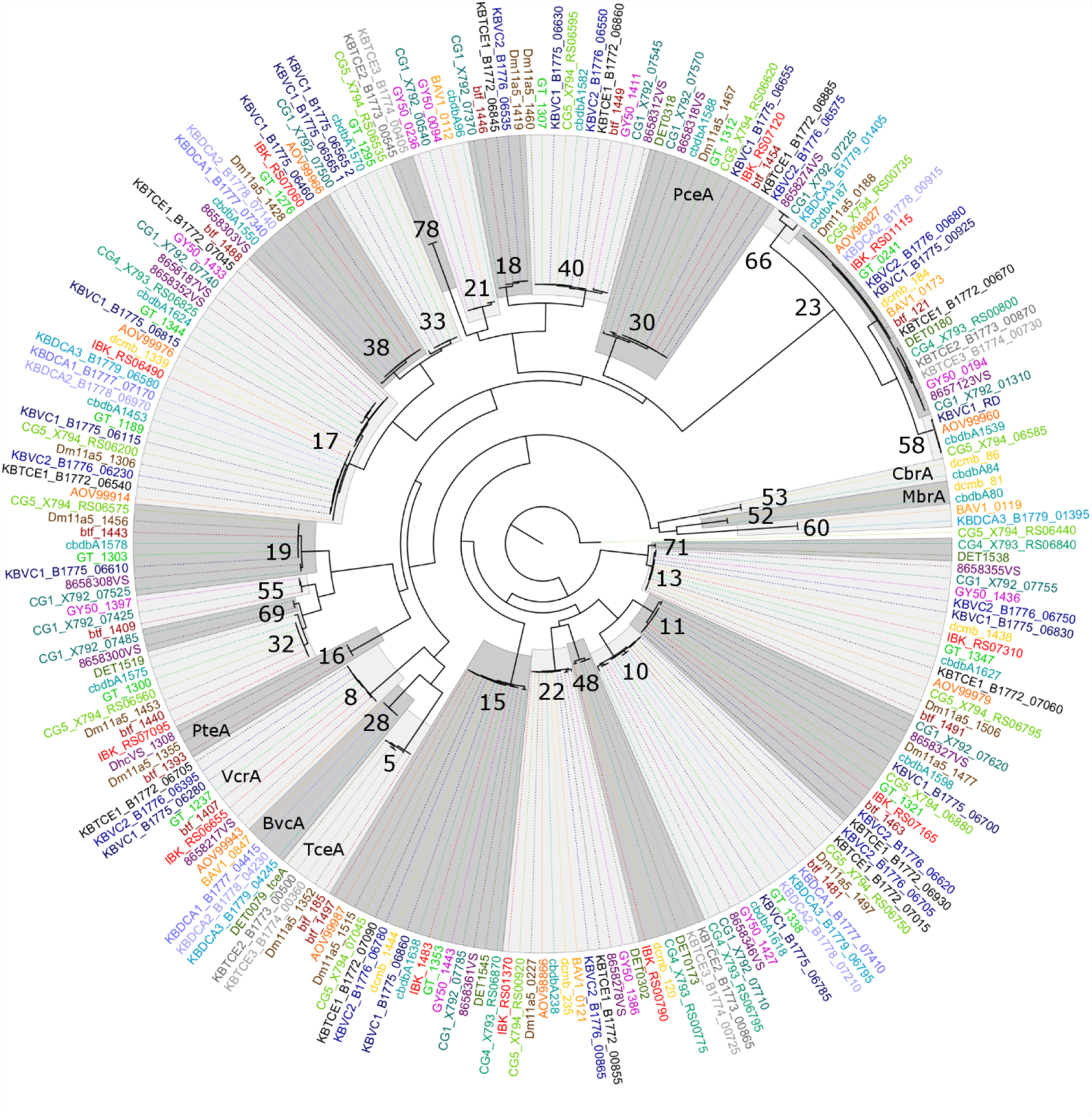
Phylogenetic amino acid tree of reductive dehalogenases from *D. mccartyi* closed genomes. Most likely tree of 100 bootstraps. Scale indicates number of substitutions per site. Orthologous groups (OGs) of dehalogenases with upwards of 90% amino acid identity are highlighted and identified by number. OGs containing a functionally characterized representative are annotated by dehalogenase name. RdhA sequences are coloured by genome they originated from. RdhA are named by NCBI locus tag. *D. mccartyi* strain name is indicated before locus tag unless included in locus tag.

Previous whole genome alignments of *D. mccartyi* have shown strong core genome region alignments (~90% nucleotide pairwise identity), with poor alignments of two regions flanking the origin, deemed the high plasticity regions (HPR) (<30% pairwise nucleotide identity) [10, 11, 87]. We identified and mapped syntenic gene blocks from KB-1 *D. mccartyi* to each other and to two Chloroflexi outgroup genomes (Figure S3). Contrary to expectation, the HPR regions were not found to be composed of unique genetic information (Figure S3). Rather, the differences between the HPRs suggested gene loss rather than recombination. When the comparison as extended to all *D. mccartyi* strains, we found a surprisingly highly conserved order of *rdhA* sequences when assigned into ortholog groups (OGs) (Figure 5A HPR1, Figure 5B HPR2, Table S2 for details). For example, at the end of HPR2 the most common order for RdhA ortholog groups is *5’*- 40-30-34-11-10-17 *-3’*. Some strains have all of these OGs (CG5, BTF08, CBDB1, KBVC1, GT, KBVC2, KBTCE1, 11a5) while certain strains only have a select few such as IBARAKI: *5’-* 30-11 *-3’*; CG4 *5’*-10-17 -*3’*; or GY50 *5’-* 40-30-10-17 *-3’* (Figure 5B, Table S2 for details).

**Figure 5A.**
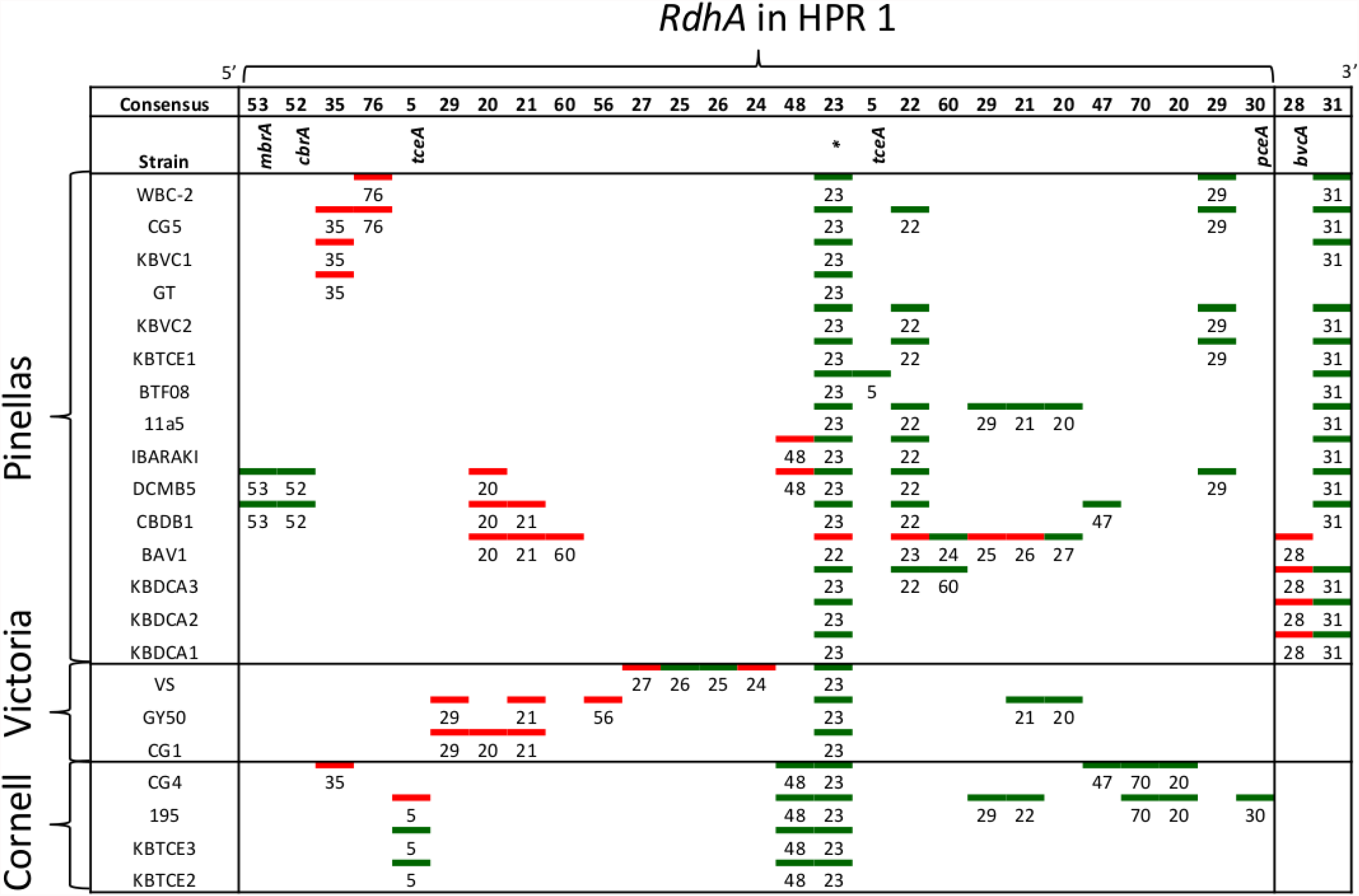
Order of *rdhA* found in high plasticity region one (HPR 1) in twenty-two *Dehalococcoides mccartyi* genomes labeled by strain name. *RdhA* are labeled by orthologous group (OG) number. *RdhA* of the same OG share >90% amino acid identity. *RdhA* without a group number are not included but can be found in Tbl.S3. Green and red accents indicate which DNA strand the gene is located on, green being leading strand clockwise from the *oriC*. The majority of *rdhA* are on the leading strand. * found in all *D. mccartyi* strains. See Table S2 for more details

**Figure 5B.**
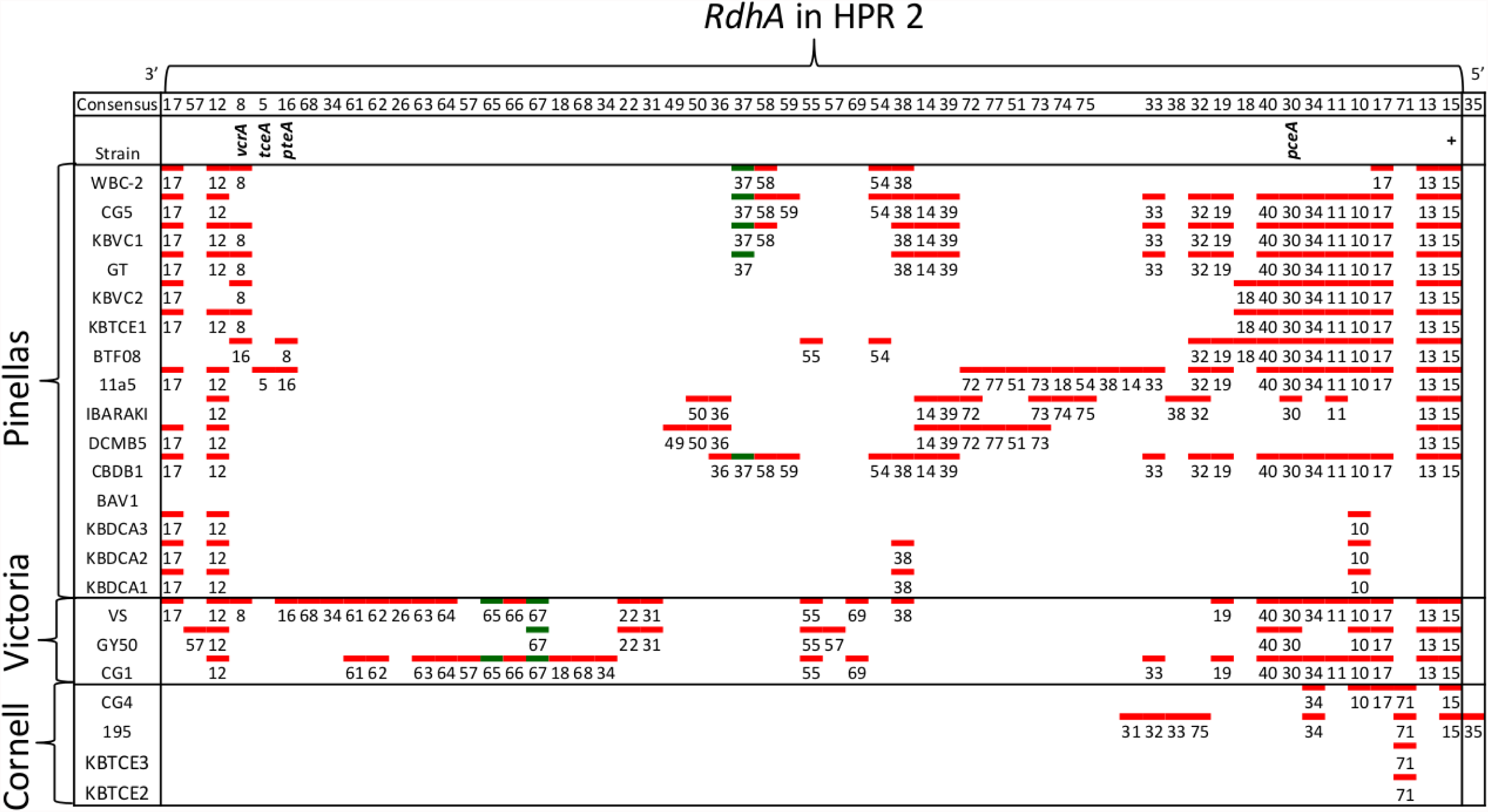
Order of *rdhA* found in high plasticity region two (HPR 2) in twenty-two *Dehalococcoides mccartyi* genomes labeled by strain name. *RdhA* are labeled by orthologous group (OG) number. *RdhA* without a group number are not included but can be found in Tbl.S1. *RdhA* of the same OG share >90% amino acid identity. Green and red accents indicate which DNA strand the gene is located on, green being leading strant clockwise from the *oriC*. The majority of *rdhA* are on the leading strand. HPR2 starts at tRNA-Leu/Arg/Val approximately 1.2 Mbp from the origin. OG 35 only in strain 195 after tRNA-Ala 1.3Mbp from the origin. + OG known to be expressed during starvation. See Table S2 for more details

### 3.4 Trends in *Dehalococcoides rdhA* acquisition, loss and evolution

For the general condition where *rdhA* occur according to the consensus order shown in Figure 5 (for more detail see Table S2), these *rdhA* are typically preceded by MarR-type regulators, such as MarR-type regulator Rdh2R (cbdb1456) found to supress downstream *rdhA* expression in *D. mccartyi* CBDB1 [88]. These *rdhA* are defined as syntenic since their location does not vary between genomes. Syntenic OG are also older than the *Dehalococcoides* clade speciation event (Figure 6). These OG contain variations in nucleotides are conserved within a particular clade which result in the clade-specific branching seen in Figure 7 (detailed examples) and S4 (larger tree). Additionally the number of nucleotide mutations observed dates their most recent common ancestral sequence earlier than when clade separation occurred (Table S3). Syntenic OG alignments have small Ka/Ks ratios suggesting that while a small fraction of mutations are deleterious, the rest are neutral meaning they are not currently under selective pressure. In summary, the first, and most common trend observed from sequence information is a consensus order of *rdhA* genes which are ancestral and not in current use (Table 2).

**Figure 6.**
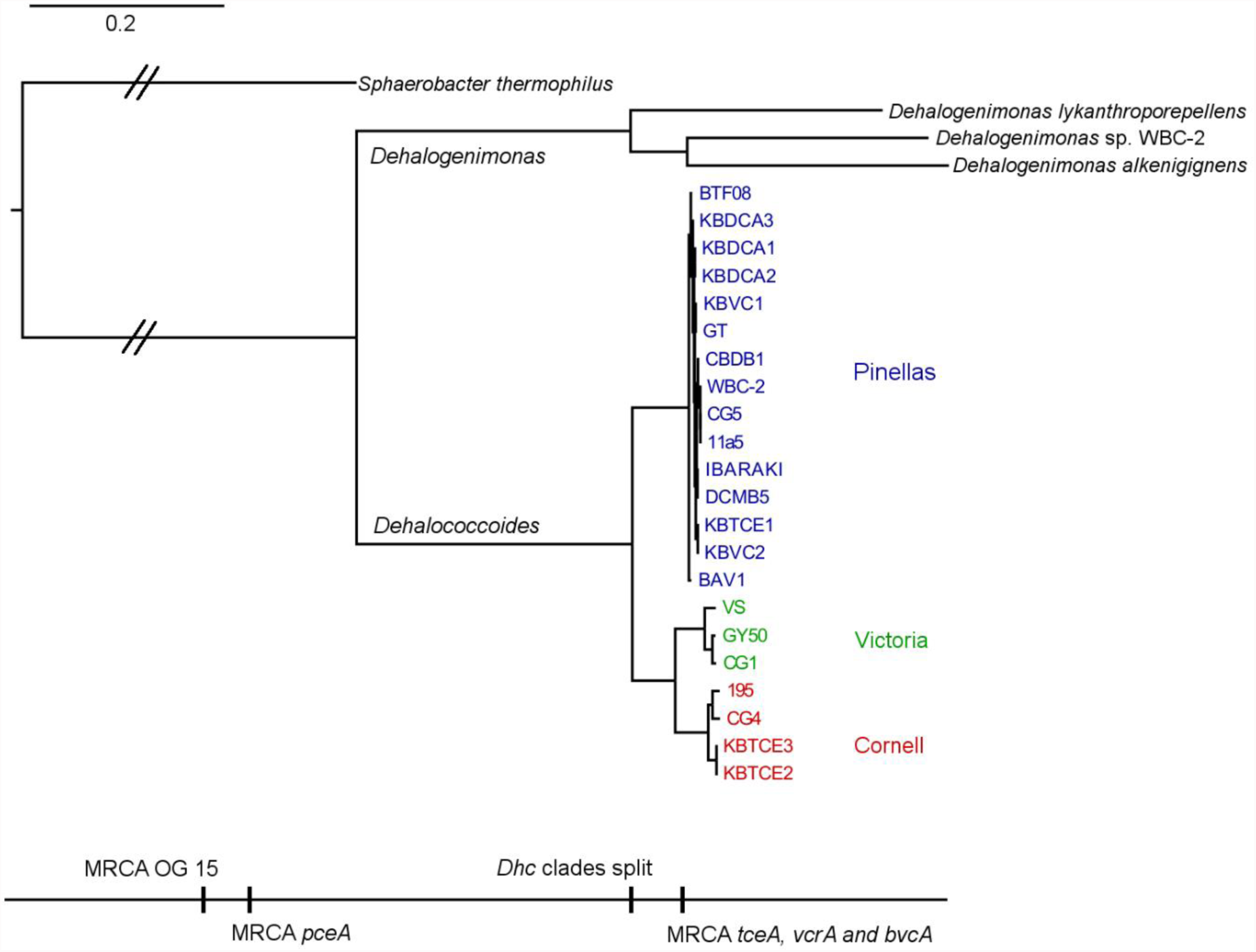
Phylogenetic tree created from an alignment of 109 concatenated core genes from *Dehalococcoides mccartyi* and *Dehalogenimonas* closed genomes with Chloroflexi *Sphaerobacter thermophilus* as out-group. Most likely tree of 100 bootstraps. Bottom scale shows timing of key events including clade separation in *D. mccartyi* and the most recent common ancestor (MRCA) of several dehalogenases listed by name or by ortholog group if uncharacterized. *D. mccartyi* clades are highlighted in common colour. Scale indicates number of substitutions per site. Double cross-hatching indicates this branch was reduced in length by half for visualization purposes.

**Figure 7.**
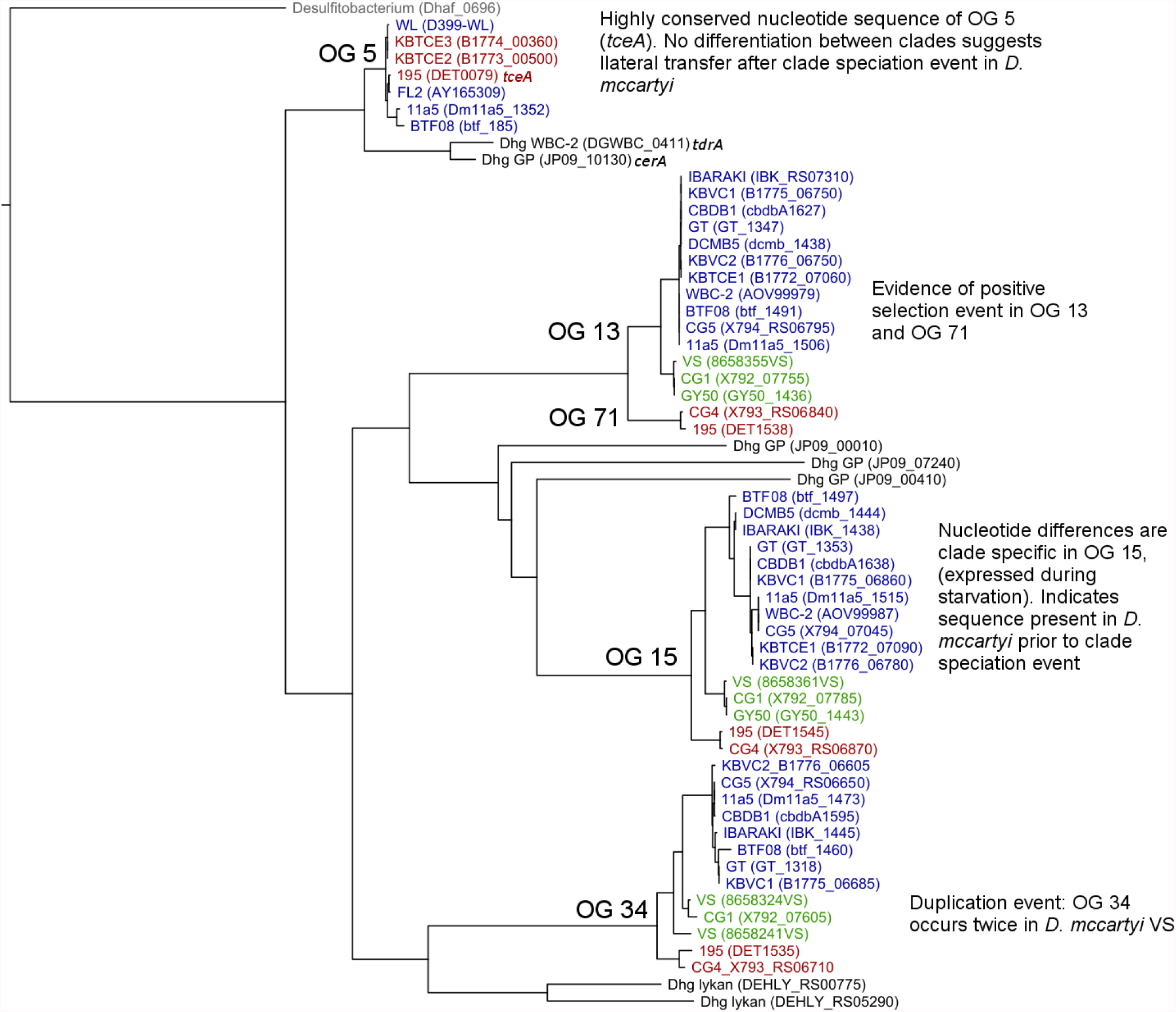
Phylogenetic tree of reductive dehalogenase genes which belong to orthologous group (OG) 5, 13, 71, 15 and 34. Most likely nucleotide tree displayed from 100 bootstraps. Scale shows number of substitutions per site. A trichloroethene dehalogenase from *Desulfitobacterium* (Dhaf_0696) used as out-group. The *rdhA* in the tree are coloured by clade: blue (Pinellas), green (Victoria), red (Cornell) and identified with strain name followed by locus tag of *rdhA* in parentheses. *Dehalogenimonas* homologous *rdhA* are shown in black listed by name if characterized in the case of *tdrA* and *cerA.*

**Table 2.** Common features of orthologous groups (OGs) of reductive dehalogenase sequences from *D. mccartyi.*

| Type | OG #s | General Features | Protein information |
| --- | --- | --- | --- |
| 1 | 10, 11, 15,<br>17, 19, 21,<br>22, 23, 30,<br>48, 55, 60,<br>66 | Present in more than one clade. Found in *D. mccartyi* before clades separated. Synteny within genomes conserved. OG 15 illustrated in Fig. 10. | OG 15 known to be expressed by *D. mccartyi* during starvation [27, 95]. OG 23 found in all *D. mccartyi* genomes at time of analysis |
| 2 | 13/71 18/40<br>32/69 33/38 | Pairs of OG groups which have recently diverged from one another. Location of *rdhA* is syntenic across all genomes. OG 13 and 71 illustrated in Fig. 10. | No characterized members at time of study |
| 3 | 5, 8, 16, 28,<br>52, 53 | Highly similar in genomes regardless of clades. Often associate with mobile genes, or on genomic island. OG 5 illustrated in Fig. 10. | 5 – TceA [30]<br>8 – VcrA [29]<br>16 – PteA [80]<br>28 – BvcA [31]<br>52 – MbrA [82]<br>53 – CbrA [96] |
| 4 | 26, 34, 57 | Duplicated within the same genome. OG 34 illustrated in Fig. 10. | No characterized members at time of study |

In four instances there is evidence for positive selection on RdhA. For example OG13 and 71 occur in the same position in the genome, but different strains present either one or the other member of the pair, but not both. OG13 (Figure 7), which is found in Pinellas and Victoria clades, is highly similar to OG71 found in the same location in the genome only in Cornell strains (Figure 7). In this case OG13 dehalogenases have small Ka/Ks ratios, while OG17 in has high ratios suggesting a positive selection of the latter. Conditions experienced by Cornell strains likely lead to the use and specialization of this dehalogenase to local conditions. We thus can define a second trend in dehalogenases in *D. mccartyi* being the evolution of new dehalogenases from existing ones (Table 2).

Three *rdhA* are thought to have been acquired horizontally by *D. mccartyi* including *tceA* (OG5, Figure 7) [17], *bvcA* (OG28) [11], and *vcrA* (OG8) [11, 16] due to their location on genomic islands and very high nucleotide and amino acid sequence conservation among strains which span vast geographical distances. Additionally, *cbrA* (OG53), *mbrA* (OG52) and *pteA* (OG16) also have very few mutations and occur in the vicinity of mobile elements. McMurdie *et al.* (2011) established that the *vcrA* gene nucleotide polymorphisms indicate that it was acquired by *Dehalococcoides* approximately ~1000 years ago, possibly earlier, after the *Dehalococcoides* clade speciation event. All *rdhA* related to mobility genes also have few mutations within OG similarly to *vcrA*, also suggesting that they were acquired in a more recent time frame (Table S3). Interestingly, all of these OGs have members which have been biochemically characterized due to their connection with industrial pollutant degradation.

Four OGs display movement within a genome, rather than between genomes (as described above). Victoria clade strains VS, CG1 and GY50 contain examples of an OG group occurring twice in the same genome suggesting a duplication event. Duplicated OGs occur in different HPRs, not in tandem (Table S2). In one case, movement of *rdhA* appears to have occurred within genome without duplication such in the case of the gene for PceA (OG30). OG 30 is syntenic in HPR2 in 14 strains with the exception of strain 195 where it occurs in HPR1 (Table S2). In strain 195, *pceA* is located near a serine recombinase which possibly mediated its movement. Additionally strain 195 *pceA* is present without its usual upstream transcriptional regulator thought to be responsible for *rdhA* regulation [88, 89]. Transcriptomic studies show that strain 195 will continue to produce high transcript levels of *pceA* regardless of starvation or TCE amendment [90]. Additionally 195 is the only strain to produce PceA in the presence of PCE. Other strains such as CG5 transcribe *pceA* in the presence of multiple PCB congeners [83] and CBDB1 was found to transcribe *pceA* in the presence of 2,3-DCP [28, 91]. Thus, the third and fourth observable trend in dehalogenase evolution is mobility, either within or between strains (Table 2).

### 3.5 Gene loss and genomic streamlining in *Dehalococcoides mccartyi*

*D. mccartyi* genomes are unique in that they are among the smallest genomes found in free-living bacteria (avg. 1.4Mbp and 1451 protein-coding genes). A common theme of all small free-living prokaryotes is their high niche specialization and low-nutrient level environments [92]. The *D. mccartyi* genome required an extensive period of time to become as specialized and small as it currently is. Wolf *et al.* 2013 [25] theorize that gene loss is equally or even more important than horizontal gene transfer in shaping genomes. The high level of synteny and number of mutations found in orthologous groups of dehalogenases uphold the same premise in *D. mccartyi*. Given that all dehalogenases whose genes show evidence of mobility are those that dehalogenate industrial contaminants, it is possible that anthropogenic releases of organohalides have caused *D. mccartyi*’s genome to enter a period of complexification by sharing select *rdhA* across vast geographic spans and causing rearrangements within genomes. The exchange of key reductive dehalogenases amongst the *D. mccartyi* is reminiscent of the recent dissemination of antibiotic resistance genes in the natural environment [93].

### 3.6 Dehalococcoidia pangenome analysis

In order to place these new genomes in relation to all of the genomes currently closed from the Dehalococcoidia, we conducted a pangenome analysis using the OrthoMCL method to cluster homologous protein groups. A total of 40,864 protein sequences from 24 *D. mccartyi* genomes and 5 *Dehalogenimonas* genomes (2 in draft) available from NCBI were used to create 2875 protein families. Of these, 623 are found in all 29 genomes representing the core-genome. The remaining 2203 protein families are part of the accessory-genome with 49 protein families being unique (i.e. only present in one strain) (Table 3 summary, all clusters available in Table S4 and S5). The first pangenome analysis conducted in 2010 from four *Dehalococcoides* genomes available at the time resulted in 1118 core genes, 457 accessory and 486 unique genes [94]. The most striking difference is that amongst the current Dehalococcoidia genomes, the number of unique protein families has been reduced from 486 to 49. A correspondence analysis (CA) was conducted on the protein families to identify the main differences between genomes (Figure 8). The CA ordination highlights the level of similarity between different strains of *Dehalococcoides mccartyi*, which could not be distinguished from one another in this analysis with statistical significance. Only the *Dehalococcoides* and the *Dehalogenimonas* genus’ were significantly different from each other (χ^2^ p-value=0.0004998, Figure S5). The *Dehalococcoides* genomes are different strains from the same species which corresponds with the outcome of the CA ordination, in contrast, the *Dehalogenimonas* genomes do come from different species, and those differences can be seen both along axis 1 (x-axis) distance from the *Dehalococcoides* cluster, and along axis 2 (y-axis) differences between the different *Dehalogenimonas* species (Figure S6 C&D). The first two dimensions accounted for 36% of the variation between the genomes (Table S6). The only protein families significantly contributing to both axis one and axis two in the CA ordination were calculated based only on the distribution of protein families in the five *Dehalogenimonas* genomes (Figure S6 B).

**Table 3.**
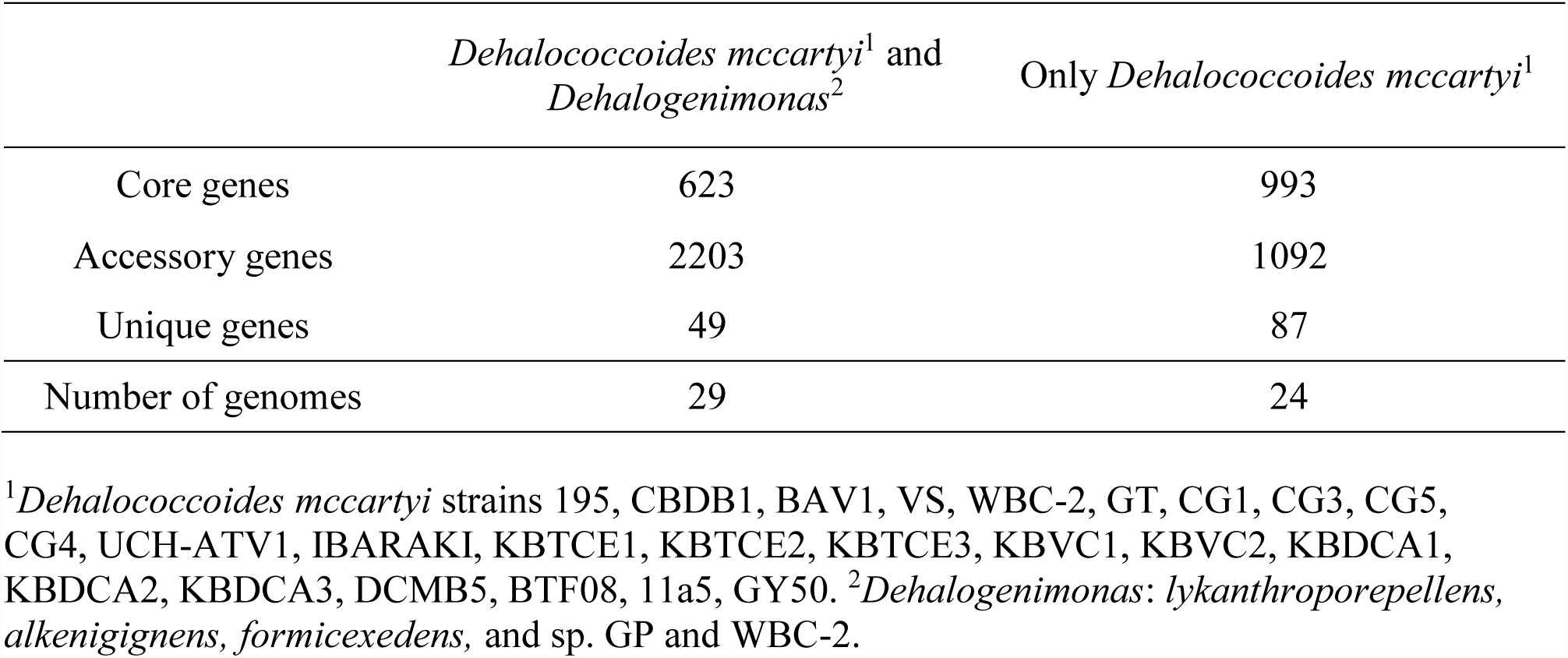
Homologous gene clustering of Dehalococcoidia pangenome from 24 *Dehalococcoides mccartyi* and 5 *Dehalogenimonas* genomes.

**Figure 8.**
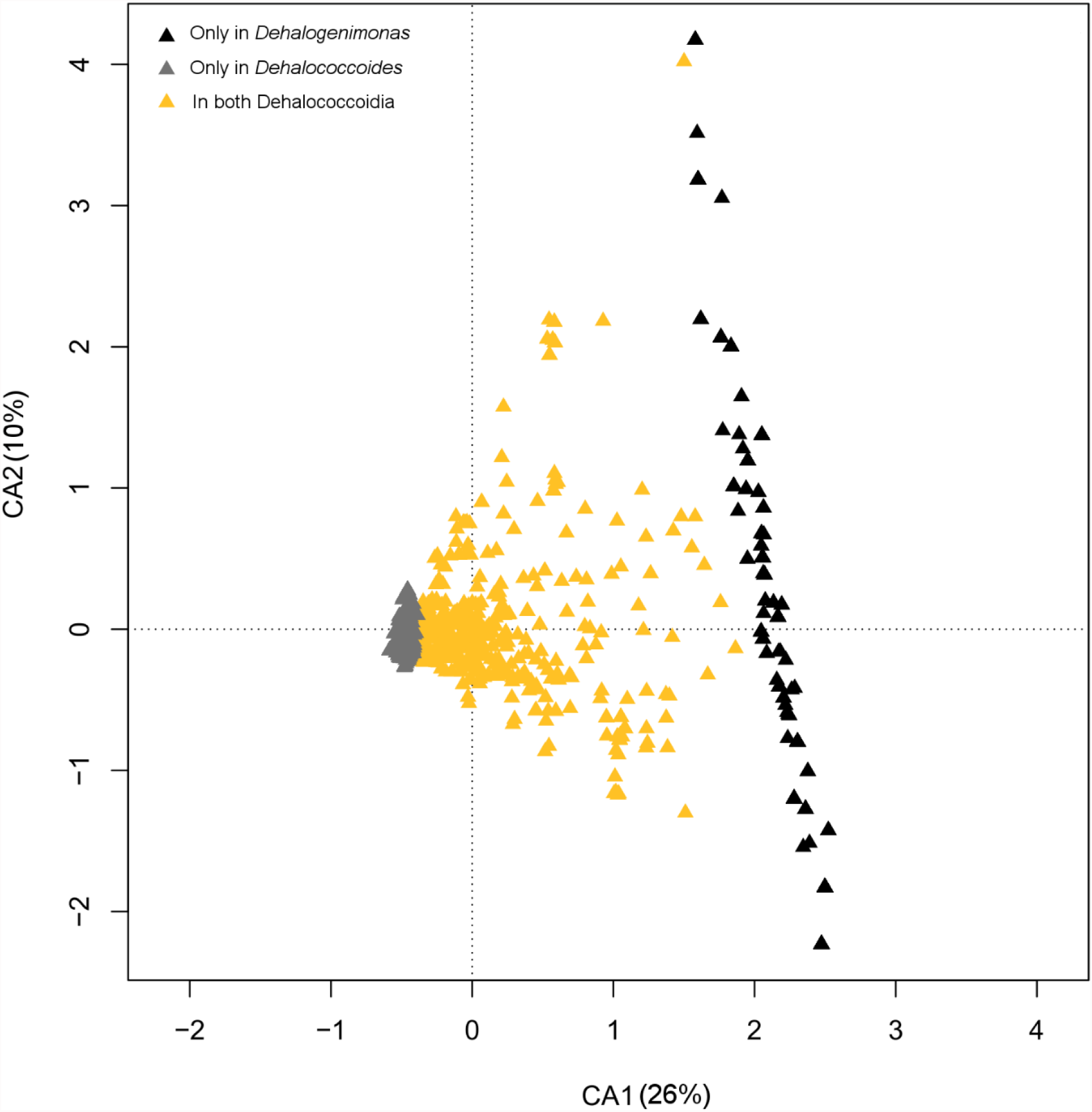
Correspondence analysis ordination plot of Dehalococcoidia pangenome. Points indicate clusters of homologous protein sequences (triangles). Clusters are coloured based on whether they contain only *Dehalococcoides* protein sequences (grey), only *Dehalogenimonas* protein sequences (black) or protein sequences from both (orange). In total 2875 clusters were identified.

Homologous protein families generated in the Dehalococcoidia pangenome analysis group all *Dehalococcoides* RdhA OGs and currently unclassified *Dehalogenimonas* RdhA into only 41 groups with roughly half (19 of 41) containing both *Dehalococcoides* and *Dehalogenimonas* RdhA (Table 4). In other words, 41 protein clusters (or families) contain RdhA sequences, and some of these clusters contain several OGs. These protein families suggest that a Dehalococcoida ancestor could have had at least 41 reductive dehalogenase genes. From all of the *rdhA* sequences found in the *Dehalococcoides* and *Dehalogenimonas* genomes it is clear that certain groups of *rdhA* are more similar to each other, and presumably have a more recent evolutionary link. When looking at *Dehalococcoides* dehalogenases alone we found that certain OGs, although still upwards of 90% conserved at the amino acid level, showed enough sequence divergence at the nucleotide level placing their most recent common ancestral gene prior to the divergence of *Dehalococcoides* and *Dehalogenimonas* (such as OG 15 Figure 7). The homologous protein clusters constructed through a pangenome analysis corroborate this premise since certain such clusters recruit sequences from the *Dehalogenimonas* along with sequences from a *Dehalococcoides-*established OG such as in the case of OG 15 (Figure 7), that is also part of homologous protein family cluster I (Table 4).

**Table 4.** Summary of contents of reductive dehalogenase (RdhA) containing homologous protein clusters. Clusters are named A to AO. Clusters which contain RdhA sequences from *Dehalococcoides* orthologous groups (OGs) are listed by group number. RdhA which have been partially biochemically characterized are listed by name.

| Cluster name | Number of Sequences | Proportion of sequences which come from *Dhc*<sup>1</sup> | OG# and characterized representatives |
| --- | --- | --- | --- |
| B | 15 | 100 | 30-PceA 70 |
| M | 19 | 100 | 13 71 |
| Q | 8 | 100 | 20 |
| T | 5 | 100 | 73 |
| X | 2 | 100 | 27 |
| Y | 4 | 100 | 50 82 |
| Z | 2 | 100 |  |
| AA | 2 | 100 | 61 |
| AB | 2 | 100 | 62 |
| AC | 3 | 100 | 80 |
| C | 31 | 86 | 18 21 37 40 |
| F | 42 | 81 | 19 32 54 55 56 57 69 |
| A | 33 | 76 | 23-conserved in all *Dhc* 58 66 72 |
| E | 85 | 73 | 17 26 33 34 36 38 63 70 75 |
| H | 23 | 69 | 5-TceA 8-VcrA 28-BvcA, 49, TdrA<sup>2</sup> CerA<sup>2</sup> |
| N | 21 | 66 | 29 47 60 65 67 |
| R | 57 | 56 | 10 11 12 24 68 |
| J | 32 | 53 | 22 39 84 81 |
| P | 5 | 45 | 53-CbrA 74 |
| O | 11 | 36 | 52-MbrA 35 |
| I | 26 | 32 | 15-expressed during starvation |
| AD | 3 | 29 | 83 |
| U | 11 | 27 | 14 |
| G | 7 | 22 | 16-PteA |
| W | 10 | 17 | 25 64 |
| AF | 2 | 17 |  |
| K | 14 | 14 | 48 |
| S | 13 | 12 | 51 59 |
| AE | 8 | 11 | 76 |
| V | 4 | 6 | DcpA<sup>2</sup> |
| D | 1 | 0 |  |
| L | 5 | 0 |  |
| AG | 2 | 0 |  |
| AH | 2 | 0 |  |
| AI | 3 | 0 |  |
| AJ | 2 | 0 |  |
| AK | 3 | 0 |  |
| AL | 4 | 0 |  |
| AM | 2 | 0 |  |
| AN | 3 | 0 |  |
| AO | 1 | 0 |  |

<sup>1</sup>Percentage of sequences which come from *Dhc* = (#*Dhc* RdhA in cluster/# *Dhc* genomes)/((#*Dhc* RdhA in cluster/# *Dhc* genomes) + (#*Dhg* RdhA in cluster/# *Dhg* genomes)) x 100.
<sup>2</sup>*Dehalogenimonas* dehalogenases

## 5.0 CONCLUSIONS

Metagenomic sequencing of the KB-1 consortium has given us new genomes and then new insights into the multiple co-existing strains of *D. mccartyi* in KB-1. The KB-1 consortium is presumed robust as a result of functional redundancy within its complement of fermenting, acetogenic, methanogenic and dechlorinationg organisms, even to the extent of including significant strain variation within the *Dehalococcoides*. *D. mccartyi* is an ancient species whose small genomes are an example of extreme genome streamlining, niche specialization and gene loss. The majority of *rdhA* genes display a much higher degree of synteny between genomes than previously appreciated and have likely been found in *D. mccartyi* for over hundreds of thousands of years, from the time of a Dehalococcoidia common ancestor. It is possible that the relatively recent anthropogenic releases of high concentrations of specific chloroorganic compounds fueled the dissemination of select reductive dehalogenases capable of their degradation, initiating a period of adaptation and complexification of the *D. mccartyi* genome. *D. mccartyi rdhA* complement has been shaped by: (1) adaptation of existing *rdhA* to new substrates (2) assimilation of new *rdhA* from the environment, or (3) within genome excision and (4) duplication or movement.

## LIST OF ABBREVIATIONS

CAH: chlorinate aliphatic hydrocarbons
PCE: perchloroethene
TCE: trichloroethene
cDCE: *cis-*dichloroethene
tDCE: *trans*-dichloroethene
VC: vinyl chloride
*rdhA*: reductive dehalogenase catalytic subunit A gene
*rdhB*: reductive dehalogenase membrane anchor gene
RdhA: reductive dehalogenase (protein) uncharacterized
RDase: reductive dehalogenase (protein)
HPR: High Plasticity Region
HPR1: High Plasticity Region 1
HPR2: High Plasticity Region 2
OG: ortholog group
SNPs: single nucleotide polymorphisms
bp: base pair
MeOH: methanol
BDBH: Bidirectional best hit
OrthoMCL: Method to identify homologous protein clusters using BDBH followed by Markov clustering

Letters A-AQ Names of homologous RdhA clusters identified by OrthoMCL

Numbers 1-81 Names of RdhA clusters (having upwards of 90% amino acid identity)

## ACKNOWLEDGEMENTS

Support was provided by the Government of Canada through Genome Canada and the Ontario Genomics Institute (no. 2009-OGI-ABC-1405 to E.A.E.), the Natural Sciences and Engineering Research Council of Canada (NSERC PGS-D to O.M., PGS B to S.T.). Support was also provided by the Government of Ontario through the ORF-GL2 program and the Unites States Department of Defense through the strategic Environmental Research and Development Program (SERDP) under contract (W912HQ-07-C-0036 project ER-1586).

## REFERENCES

1. Stroo HF, Leeson A, Ward CH: Bioaugmentation for Groundwater Remediation: Springer New York; 2012.

2. Major DJ, McMaster ML, Cox EE, Edwards EA, Dwortzek SM, Hendrickson ER, Starr MG, Payne JA, Buonamici LW: Field demonstration of sucessful bioaugmentation to achieve dechlorination of tetrachloroethene to ethene. Environ Sci Technol 2000, 36(23):5106–5116.

3. Duhamel M, S. D. Wehr, L. Yu, H. Rizvi, D. Seepersad, S. Dworatzek, E. E. Cox, and E. A. Edwards.: Comparison of anaerobic dechlorinating enrichment cultures maintained on tetrachloroethene, trichloroethene, *cis*-dichloroethene and vinyl chloride. Water Res 2002:36,4193–4202.

4. Maymo-Gatell X, Chien Y, Gossett JM, Zinder SH: Isolation of a bacterium that reductively dechlorinates tetrachloroethene to ethene. Science 1997, 276:1568–1571.

5. He J, Ritalahti KM, Yang KL, Koenigsberg SS, Löffler FE: Detoxification of vinyl chloride to ethene coupled to growth of an anaerobic bacterium. Nature 2003, 424(6944):62–65.

6. Yang Y, Higgins SA, Yan J, Simsir B, Chourey K, Iyer R, Hettich RL, Baldwin B, Ogles DM, Löffler FE: Grape pomace compost harbors organohalide-respiring *Dehalogenimonas* species with novel reductive dehalogenase genes. ISME J 2017.

7. Molenda O, Quaile AT, Edwards EA: *Dehalogenimonas* sp. strain WBC-2 genome and identification of its *trans*-dichloroethene reductive dehalogenase, TdrA. Appl Environ Microbiol 2015, 82(1):40–50.

8. Duhamel M, Wehr SD, Yu L, Rizvi H, Seepersad D, Dworatzek S, Cox EE, Edwards EA: Comparison of anaerobic dechlorinating enrichment cultures maintained on tetrachloroethene, trichloroethene, *cis*-dichloroethene and vinyl chloride. Water Res 2002, 36(17):4193–4202.

9. Calabrese EJ, Kostecki PT, Dragun J: Contaminated Soils, Sediments and Water: Science in the Real World: Springer US; 2006.

10. Kube M, Beck A, Zinder SH, Kuhl H, Reinhardt R, Adrian L: Genome sequence of the chlorinated compound-respiring bacterium *Dehalococcoides* species strain CBDB1. Nat Biotech 2005, 23(10):1269–1273.

11. McMurdie PJ, Behrens SF, Müller JA, Goke J, Ritalahti KM, Wagner R, Goltsman E, Lapidus A, Holmes S, Löffler FE et al: Localized plasticity in the streamlined genomes of vinyl chloride respiring *Dehalococcoides*. PLoS Genet 2009, 5(11):e1000714.

12. Grindley ND, Whiteson KL, Rice PA: Mechanisms of site-specific recombination. Annu Rev Biochem 2006, 75:567–605.

13. Komano T, Kim SR, Yoshida T, Nisioka T: DNA rearrangement of the shufflon determines recipient specificity in liquid mating of IncI1 plasmid R64. J Mol Biol 1994, 243(1):6–9.

14. Johnson RC: Site-specific DNA Inversion by Serine Recombinases. Microbiology spectrum 2015, 3(1):Mdna3–0047-2014.

15. Birge EA: Site-specific recombination following conjugation in *Escherichia coli* K-12. Mol Gen Genet 1983, 192(3):366–372.

16. McMurdie PJ, Hug LA, Edwards EA, Holmes S, Spormann AA,: Site-specific mobilization of vinyl-chloride respiration islands by a mechanism common in *Dehalococcoides*. BMC Genomics 2011, 12:287–302.

17. Krajmalnik-Brown R, Sung Y, Ritalahti KM, Saunders FM, Löffler FE: Environmental distribution of the trichloroethene reductive dehalogenase gene (*tceA*) suggests lateral gene transfer among *Dehalococcoides*. FEMS Microbiol Ecol 2007, 59(1):206–214.

18. Hug LA, Maphosa F, Leys D, Löffler FE, Smidt H, Edwards EA, Adrian L: Overview of organohalide-respiring bacteria and a proposal for a classification system for reductive dehalogenases. Philos Trans R Soc Biol Sci 2013, 368:1–10.

19. Krzmarzick MJ, Crary BB, Harding JJ, Oyerinde OO, Leri AC, Myneni SC, Novak PJ: Natural niche for organohalide-respiring Chloroflexi. Appl Environ Microbiol 2012, 78(2):393–401.

20. Martínez-Cano DJ, Reyes-Prieto M, Martínez-Romero E, Partida-Martínez LP, Latorre A, Moya A, Delaye L: Evolution of small prokaryotic genomes. Front Microbiol 2014, 5:742.

21. Dufresne A, Garczarek L, Partensky F: Accelerated evolution associated with genome reduction in a free-living prokaryote. Genome biology 2005, 6(2):R14.

22. Mira A, Ochman H, Moran NA: Deletional bias and the evolution of bacterial genomes. Trends Genet 2001, 17(10):589–596.

23. Marais GA, Calteau A, Tenaillon O: Mutation rate and genome reduction in endosymbiotic and free-living bacteria. Genetica 2008, 134(2):205–210.

24. Berg OG, Kurland CG: Evolution of microbial genomes: sequence acquisition and loss. Mol Biol Evol 2002, 19(12):2265–2276.

25. Wolf YI, Koonin EV: Genome reduction as the dominant mode of evolution. Bioessays 2013, 35(9):829–837.

26. Hölscher T, Krajmalnik-Brown R, Ritalahti KM, von Wintzingerode F, Görisch H, Löffler FE, Adrian L: Multiple nonidentical reductive-dehalogenase-homologous genes are common in *Dehalococcoides*. Appl Environ Microbiol 2004, 70(9):5290–5297.

27. Liang X, Molenda O, Tang S, Edwards EA: Identity and substrate-specificity of reductive dehalogenases expressed in *Dehalococcoides-*containing enrichment cultures maintained on different chlorinated ethenes. Appl Environ Microbiol 2015, 81(14):4626–4633.

28. Fung JM, Morris RM, Adrian L, Zinder SH: Expression of reductive dehalogenase genes in *Dehalococcoides ethenogenes* strain 195 growing on tetrachloroethene, trichloroethene, or 2,3-dichlorophenol. Appl Environ Microbiol 2007, 73(14):4439–4445.

29. Müller JA, Rosner BM, Von Abendroth G, Meshulam-Simon G, McCarty PL, Spormann AM: Molecular identification of the catabolic vinyl chloride reductase from *Dehalococcodies* sp. strain VS and its environmental distribution. Appl Environ Microbiol 2004, 70(8):4880–4888.

30. Magnuson JK, Romine MF, Burris DR, Kingsley MT: Trichloroethene reductive dehalogenase from *Dehalococcoides ethenogenes:* sequence of *tceA* and substrate range characterization. Appl Environ Microbiol 2000, 66(12):5141–5147.

31. Tang S, Chan WMW, Fletcher KE, Seifert J, Liang X, Löffler FE, Edwards EA, Adrian L: Functional characterizayion of reductive dehalogenases by using blue native polyacrylamide gel electrophoresis. Appl Environ Microbiol 2013, 79(3):974–981.

32. Chow WL, Cheng D, Wang S, He J: Identification and transcription analysis of *trans-*DCE producing reductive dehalogenases in *Dehalococcoides* species. The ISME Journal 2010, 4:1020–2013.

33. Adrian L, Rahnenführer J, Gobom J, Hölscher T: Identification of a Chlorobenzene Reductive Dehalogenase in *Dehalococcoides* sp. Strain CBDB1. Appl Environ Microbiol 2007, 73(23):7717–7724.

34. Zhao S, Ding C, He J: Genomic characterization of *Dehalococcoides mccartyi* strain 11a5 reveals a circular extrachromosomal genetic element and a new tetrachloroethene reductive dehalogenase gene. FEMS Microbiol Ecol 2017, 93(4):fiw235–fiw235.

35. Molenda O, Quaile AT, Edwards EA: *Dehalogenimonas* sp. strain WBC-2 genome and identification of its *trans*-dichloroethene reductive dehalogenase, TdrA. Appl Environ Microbiol 2016, 82(1):40–50.

36. Padilla-Crespo E, Yan J, Swift C, Wagner DD, Chourey K, Hettich RL, Ritalahti KM, Löffler FE: Identification and environmental distribution of *dcpA*, which encodes the reductive dehalogenase catalyzing the dichloroelimination of 1,2-dichloropropane to propene in organo-halide-respiring chloroflexi. Appl Environ Microbiol 2014, 80(3):808–818.

37. Maphosa F, de Vos WM, Smidt H: Exploiting the ecogenomics toolbox for environmental diagnostics of organohalide-respiring bacteria. Trends Biotechnol 2010, 28(6):308–316.

38. Bommer M, Kunze C, Fesseler J, Schubert T, Diekert G, Dobbek H: Structural basis for organohalide respiration. Science 2014, 346(6208):455–458.

39. Payne KA, Quezada CP, Fisher K, Dunstan MS, Collins FA, Sjuts H, Levy C, Hay S, Rigby SE, Leys D: Reductive dehalogenase structure suggests a mechanism for B12- dependent dehalogenation. Nature 2015, 517(7535):513–516.

40. Maymo-Gatell X, Chien Y, Gossett JM, Zinder SH: Isolation of a bacterium that reductively dechlorinates tetrachloroethene to ethene. Science 1997, 276(5318):1568–1571.

41. Sung Y, Ritalahti KM, Apkarian RP, Löffler FE: Quantitative PCR confirms purity of strain GT, a novel trichloroethene-to-ethene-respiring *Dehalococcoides* isolate. Appl Environ Microbiol 2006, 72(3):1980–1987.

42. Duhamel M, Mo K, Edwards EA: Characterization of a highly enriched *Dehalococcoides*-containing culture that grows on vinyl chloride and trichloroethene. Appl Environ Microbiol 2004, 70(9):5538–5545.

43. Pérez-de-Mora A, Lacourt A, McMaster ML, Liang X, Dworatzek SM, Edwards EA: Chlorinated electron acceptor abundance drives selection of *Dehalococcoides mccartyi* (*D. mccartyi*) strains in dechlorinating enrichment cultures and groundwater environments. Front Microbiol 2018, 9(812).

44. Tang S, Gong Y, Edwards EA: Semi-automatic in silico gap closure enabled de novo assembly of two *Dehalobacter* genomes from metagenomic data. PLoS One 2012, 7(12).

45. Bolger AM, Lohse M, Usadel B: Trimmomatic: a flexible trimmer for Illumina sequence data. Bioinformatics 2014, 30(15):2114–2120.

46. Simpson JT, Wong K, Jackman SD, Schein JE, Jones SJ, Birol I: ABySS: a parallel assembler for short read sequence data. Genome Res 2009, 19(6):1117–1123.

47. Boetzer M, Henkel CV, Jansen HJ, Butler D, Pirovano W: Scaffolding pre-assembled contigs using SSPACE. Bioinformatics 2011, 27(4):578–579.

48. Gnerre S, Maccallum I, Przybylski D, Ribeiro FJ, Burton JN, Walker BJ, Sharpe T, Hall G, Shea TP, Sykes S et al: High-quality draft assemblies of mammalian genomes from massively parallel sequence data. Proc Natl Acad Sci U S A 2011, 108(4):1513–1518.

49. Brown CT, Howe A, Zhang Q, Pyrokosz AB, Brom TH: A reference-free algorithm for computational normalization of shotgun sequencing data. Quantitative Biology 2012.

50. Kearse M, Moir R, Wilson A, Stones-Havas S, Cheung M, Sturrock S, Buxton S, Cooper A, Markowitz S, Duran C et al: Geneious Basic: an integrated and extendable desktop software platform for the organization and analysis of sequence data. Bioinformatics 2012, 28(12):1647–1649.

51. Aziz R, Bartels D, Best AD, M, Disz T, Edwards R, Forsma K, Gerdes SG, EM, Kubal, M, Meyer F, Olsen G et al: The RAST server: rapid annotations using subsystems technology. BMC Genomics 2008, 9(75).

52. Van Domselaar GH, Stothard P, Shrivastava S, Cruz JA, Guo A, Dong X, Lu P, Szafron D, Greiner R, Wishart DS: BASys: a web server for automated bacterial genome annotation. Nucleic Acids Res 2005, 33(Web Server issue):W455–W459.

53. Tatusova T, DiCuccio M, Badretdin A, Chetvernin V, Nawrocki EP, Zaslavsky L, Lomsadze A, Pruitt KD, Borodovsky M, Ostell J: NCBI prokaryotic genome annotation pipeline. Nucleic Acids Res 2016, 44(14):6614–6624.

54. Frank A, Lobry J: Oriloc: prediction of replication boundaries in unannotated bacterial chromosomes. Bioinformatics 2000, 16:566–567.

55. Engelbrektson A, Kunin V, Wrighton KC, Zvenigorodsky N, Chen F, Ochman H, Hugenholtz P: Experimental factors affecting PCR-based estimates of microbial species richness and evenness. ISME J 2010, 4(5):642–647.

56. Ramos-Padron E, Bordenave S, Lin S, Bhaskar IM, Dong X, Sensen CW, Fournier J, Voordouw G, Gieg LM: Carbon and sulfur cycling by microbial communities in a gypsum-treated oil sands tailings pond. Environ Sci Technol 2011, 45(2):439–446.

57. Caporaso JG, Kuczynski J, Stombaugh J, Bittinger K, Bushman FD, Costello EK, Fierer N, Peña AG, Goodrich JK, Gordon JI et al: QIIME allows analysis of high-throughput community sequencing data. Nat Methods 2010, 7(5).

58. Edgar RC: Search and clustering orders of magnitude faster than BLAST. Bioinformatics 2010, 26(19):2460–2461.

59. DeSantis TZ, Hugenholtz P, Larsen N, Rojas M, Brodie EL, Keller K, Huber T, Dalevi D, Hu P, Andersen GL: Greengenes, a chimera-checked 16S rRNA gene database and workbench compatible with ARB. Appl Environ Microbiol 2006, 72(7):5069–5072.

60. Wang Q, Garrity GM, Tiedje JM, Cole JR: Naive Bayesian classifier for rapid assignment of rRNA sequences into the new bacterial taxonomy. Appl Environ Microbiol 2007, 73(16):5261–5267.

61. Grostern A, Edwards EA: Characterization of a *Dehalobacter* coculture that dechlorinates 1,2-dichloroethane to ethene and identification of the putative reductive dehalogenase gene. Appl Environ Microbiol 2009, 75(9):2684–2693.

62. Waller AS, Krajmalnik-Brown R, Löffler FE, Edwards EA: Multiple reductive-dehalogenase-homologous genes are simultaneously transcribed during dechlorination by *Dehalococcoides*-containing cultures. Appl Environ Microbiol 2005, 71(12):8257–8264.

63. Ritalahti KM, Amos BK, Sung Y, Wu Q, Koenigsberg SS, Löffler FE: Quantitative PCR targeting 16S rRNA and multiple reductive dehalogenase genes simultaneously monitors multiple *Dehalococcoides* strains. Appl Environ Microbiol 2006, 72(4):2765–2774.

64. Edgar RC: MUSCLE: multiple sequence alignment with high accuracy and high throughput. Nucleic Acids Res 2004, 32(5):1792–1797.

65. Stamatakis A: RAxML-VI-HPC: maximum likelihood-based phylogenetic analyses with thousands of taxa and mixed models. Bioinformatics 2006, 22(21):2688–2690.

66. Ahn YB, Kerkhof LJ, Haggblom MM: *Desulfoluna spongiiphila* sp. nov., a dehalogenating bacterium in the Desulfobacteraceae from the marine sponge Aplysina aerophoba. Int J Syst Evol Microbiol 2009, 59(Pt 9):2133–2139.

67. Nekrutenko A, Makova KD, Li W-H: The K(A)/K(S) Ratio Test for Assessing the Protein-Coding Potential of Genomic Regions: An Empirical and Simulation Study. Genome Res 2002, 12(1):198–202.

68. Contreras-Moreira B, Vinuesa P: GET_HOMOLOGUES, a versatile software package for scalable and robust microbial pangenome analysis. Appl Environ Microbiol 2013, 79(24):7696–7701.

69. Darling AE, Mau B, Perna NT: progressiveMauve: multiple genome alignment with gene gain, loss and rearrangement. PLoS One 2010, 5(6):e11147.

70. Wang Y, Tang H, Debarry JD, Tan X, Li J, Wang X, Lee TH, Jin H, Marler B, Guo H et al: MCScanX: a toolkit for detection and evolutionary analysis of gene synteny and collinearity. Nucleic Acids Res 2012, 40(7):e49.

71. Hug LA, Beiko RG, Rowe AR, Richardson RE, Edwards EA: Comparative metagenomics of three *Dehalococcoides*-containing enrichment cultures: the role of the non-dechlorinating community. BMC Genomics 2012, 13:327–327.

72. Waller A: Molecular investigation of chloroethene reductive dehalogenation by the mixed microbial community KB-1. University of Toronto; 2010.

73. Waller AS, Hug LA, Mo K, Radford DR, Maxwell KL, Edwards EA: Transcriptional analysis of a *Dehalococcoides*-containing microbial consortium reveals prophage activation. Appl Environ Microbiol 2012, 78(4):1178–1186.

74. Molenda O, Tang S, Edwards EA: Complete genome sequence of *Dehalococcoides mccartyi* strain WBC-2, capable of anaerobic reductive dechlorination of vinyl chloride. Genome Announc 2016, 4(6):e01375–01316.

75. Jones EJP, Voytek MA, Lorah MM, Kirshtein JD: Characterization of a microbial consortium capable of rapid and simultaneous dechlorination of 1,1,2,2- tetrachloroethane and chlorinated ethane and ethene intermediates. Bioremediation J 2006, 10(4):153–168.

76. Lorah MM, Majcher EH, Jones EJ, Voytek MA: Microbial consortia development and microcosm and column experiments for enhanced bioremediation of chlorinated volatile organic compounds, West Branch Canal Creek Wetland Area, Aberdeen Proving Ground, Maryland. In. Edited by Survey UDotIUG. Reston, Virginia: US Scientific Investigations Report 2007-5165; 2007.

77. Löffler FE, Yan J, Ritalahti KM, Adrian L, Edwards EA, Konstantinidis KT, Müller JA, Fullerton H, Zinder SH, Spormann AM: *Dehalococcoides mccartyi* gen. nov., sp. nov., obligately organohalide-respiring anaerobic bacteria relevant to halogen cycling and bioremediation, belong to a novel bacterial class, *Dehalococcoidia* classis nov., order *Dehalococcoidales* ord. nov. and family *Dehalococcoidaceae* fam. nov., within the phylum Chloroflexi. Int J Syst Evol Microbiol 2013, 63(Pt 2):625–635.

78. Hendrickson ER, Payne JA, Young RM, Starr MG, Perry MP, Fahnestock S, Ellis DE, Ebersole RC: Molecular Analysis of *Dehalococcoides* 16S Ribosomal DNA from Chloroethene-Contaminated Sites throughout North America and Europe. Appl Environ Microbiol 2002, 68(2):485–495.

79. Pöritz M, Goris T, Wubet T, Tarkka MT, Buscot F, Nijenhuis I, Lechner U, Adrian L: Genome sequences of two dehalogenation specialists - *Dehalococcoides* mccartyi strains BTF08 and DCMB5 enriched from the highly polluted Bitterfeld region. FEMS Microbiol Lett 2013, 343(2):101–104.

80. Zhao S, Ding C, He J: Genomic characterization of *Dehalococcoides mccartyi* strain 11a5 reveals a circular extrachromosomal genetic element and a new tetrachloroethene reductive dehalogenase gene. FEMS Microbiol Ecol 2016.

81. Yohda M, Yagi O, Takechi A, Kitajima M, Matsuda H, Miyamura N, Aizawa T, Nakajima M, Sunairi M, Daiba A et al: Genome sequence determination and metagenomic characterization of a *Dehalococcoide*s mixed culture grown on *cis*-1,2- dichloroethene. J Biosci Bioeng 2015, 120(1):69–77.

82. Cheng D, He J: Isolation and characterization of “*Dehalococcoides*” sp. strain MB which dechlorinates tetrachloroethene to *trans-*1,2-dichloroethene. Appl Environ Microbiol 2009, 75(18):5910–5918.

83. Wang S, Chng KR, Wilm A, Zhao S, Yang K-L, Nagarajan N, He J: Genomic characterization of three unique *Dehalococcoides* that respire on persistent polychlorinated biphenyls. PNAS 2014, 111(33):12103–12108.

84. Molenda O, Tang S, Lomheim L, Vasu G, Lemak S, Yakunin AF, Edwards EA: Extrachromosomal circular elements targeted by CRISPR-Cas in *Dehalococcoides mccartyi* are linked to mobilization of reductive dehalogenase genes. ISME J 2018, *accepted*.

85. Low A, Shen Z, Cheng D, Rogers MJ, Lee PKH, He J: A comparative genomics and reductive dehalogenase gene transcription study of two chloroethene-respiring bacteria, *Dehalococcoides mccartyi* strains MB and 11a. Scientific Reports 2015, 5:15204.

86. Lee PKH, He J, Zinder SH, Alvarez-Cohen L: Evidence for nitrogen fixation by “*Dehalococcoides ethenogenes*” strain 195. Appl Environ Microbiol 2009, 75(23):7551–7555.

87. Seshardi R, Adrian L, Fouts DE, Eisen JA, Phillippy AM, Methe BA, Ward NL, Nelson WC, Dobson RJ, Daugherty SC et al: Genome sequence of the PCE-dechlorinating bacterium *Dehalococcoides ethenogenes*. Science 2005 307(5706):105–108.

88. Wagner A, Segler L, Kleinsteuber S, Sawers G, Smidt H, Lechner U: Regulation of reductive dehalogenase gene transcription in *Dehalococcoides mccartyi*. Philos Trans R Soc Biol Sci 2013, 368(1616):20120317.

89. Krasper L, Lilie H, Kublik A, Adrian L, Golbik R, Lechner U: The MarR-Type Regulator Rdh2R Regulates *rdh* Gene Transcription in *Dehalococcoides mccartyi* strain CBDB1. J Bacteriol 2016, 198(23):3130–3141.

90. Mansfeldt CB, Rowe AR, Heavner GLW, Zinder SH, Richardson RE: Meta-analyses of *Dehalococcoides mccartyi* strain 195 transcriptomic profiles identify a respiration rate-related gene expression transition point and interoperon recruitment of a key oxidoreductase subunit. Appl Environ Microbiol 2014, 80(19):6062–6072.

91. Morris RM, Fung JM, Rahm BG, Zhang S, Freedman DL, Zinder SH, Richardson RE: Comparative proteomics of *Dehalococcoides* spp. reveals strain-specific peptides associated with activity. Appl Environ Microbiol 2007, 73(1):320–326.

92. Lauro FM, McDougald D, Thomas T, Williams TJ, Egan S, Rice S, DeMaere MZ, Ting L, Ertan H, Johnson J et al: The genomic basis of trophic strategy in marine bacteria. Proc Natl Acad Sci U S A 2009, 106(37):15527–15533.

93. Allen HK, Donato J, Wang HH, Cloud-Hansen KA, Davies J, Handelsman J: Call of the wild: antibiotic resistance genes in natural environments. Nat Rev Micro 2010, 8(4):251–259.

94. Islam MA, Waller AS, Hug LA, Provart NJ, Edwards EA, Mahadevan R: New Insights into *Dehalococcoides mccartyi* Metabolism from a Reconstructed Metabolic Network-Based Systems-Level Analysis of *D. mccartyi* Transcriptomes. PLoS ONE 2014, 9(4):e94808.

95. Johnson DR, Brodie EL, Hubbard AE, Andersen GL, Zinder SH, Alvarez-Cohen L: Temporal Transcriptomic Microarray Analysis of “Dehalococcoides ethenogenes” Strain 195 during the Transition into Stationary Phase. Applied and Environmental Microbiology 2008, 74(9):2864–2872.

96. Adrian L, Rahnenfuhrer J, Gobom J, Holscher T: Identification of a chlorobenzene reductive dehalogenase in Dehalococcoides sp. strain CBDB1. Appl Environ Microbiol 2007, 73(23):7717–7724.

